# Systems engineering of *Saccharomyces cerevisiae* for synthesis and accumulation of vanillin

**DOI:** 10.1101/2023.06.16.545388

**Authors:** Qiwen Mo, Jifeng Yuan

## Abstract

Vanillin represents one of the most widely used flavoring agents in the world. However, microbial synthesis of vanillin is hindered by the host native metabolism that could rapidly degrade vanillin to the byproducts. Here, we report that the industrial workhorse *Saccharomyces cerevisiae* was engineered by systematic deletion of oxidoreductases to improve the vanillin accumulation. Subsequently, we harnessed the **r**educed aromatic **a**ldehyde **re**duction (RARE) yeast platform for *de novo* synthesis of vanillin from glucose. We investigated multiple coenzyme-A free pathways to improve vanillin production in yeast. The vanillin productivity in yeast was further enhanced by systems engineering to optimize the supply of cofactors (NADPH and *S*-adenosylmethionine) together with metabolic reconfiguration of yeast central metabolism. The final yeast strain with overall 24 genetic modifications produced 365.55 ± 7.42 mg L^−1^ under shake-flasks, which represents the highest vanillin titer from glucose achieved to date. Taken together, our work lays a foundation for the future implementation of vanillin production from glucose in budding yeast.

TOC

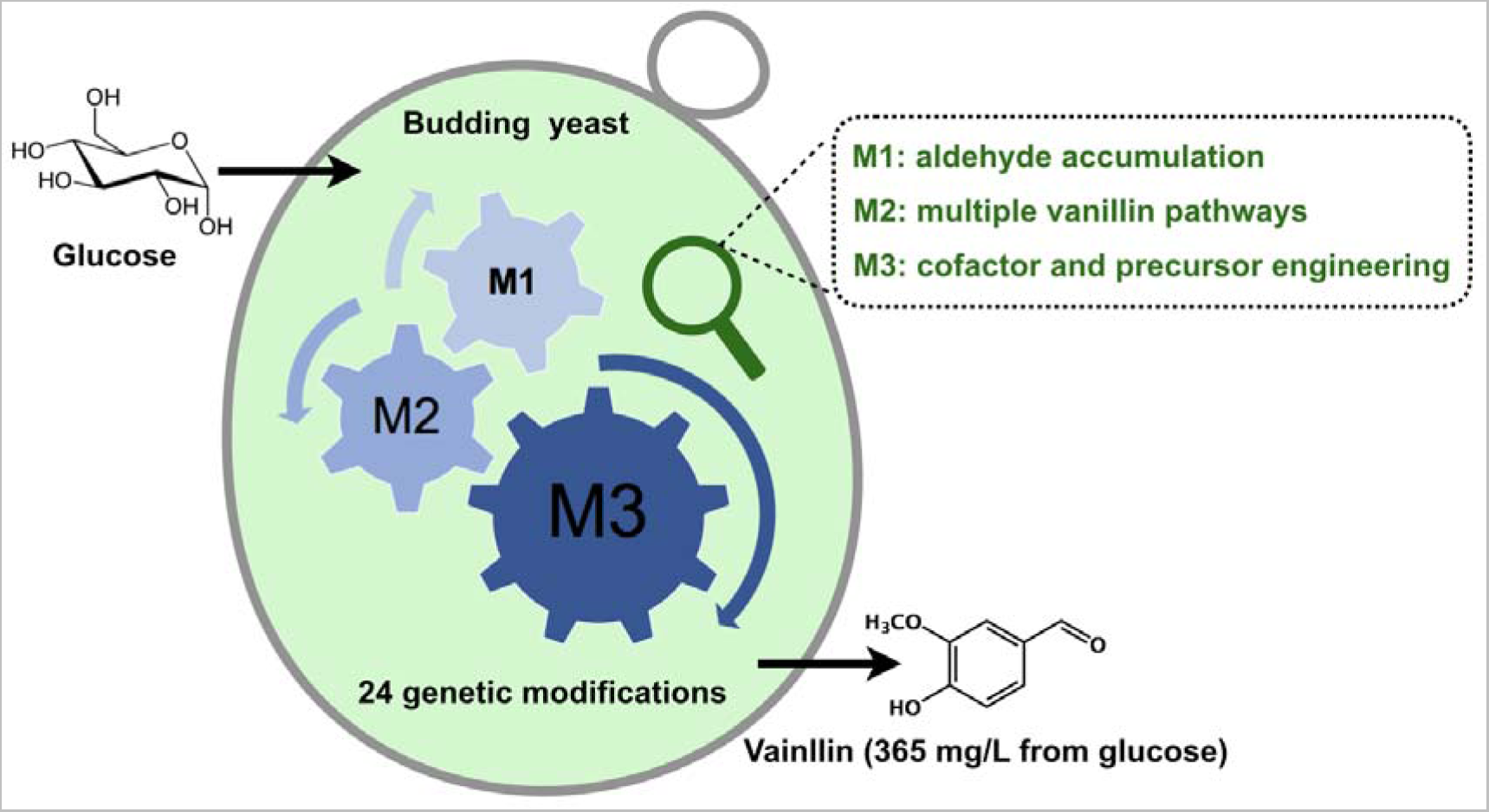

## Introduction

Aromatic aldehydes are important flavor and fragrance compounds. For instance, vanillin has an intense and tenacious creamy vanilla-like taste, which makes it one of the most widely used flavoring agents in the world ^1^. It is estimated that more than 16,000 tons of vanillin are consumed annually across the world. In addition, vanillin possesses antimicrobial, antioxidant, antimutagenic, hypolipidemic, anti-sickling, and anti-inflammatory activities ^2–4^. It is also a crucial raw ingredient in the manufacture of pharmaceutical medications such as dopamine and Aldomet ^5^. As a plant secondary metabolite, vanillin can be extracted from the seedpods of orchids such as *Vanilla*. Because of the vanilla orchid’s sluggish development and very low vanillin concentration in the mature vanilla pods, the plant-sourced vanillin comes with a relatively high cost (US$ 515 kg^−1^ in June 2018) ^1, 6^. Although vanillin can be synthesized from fossil hydrocarbons with a cheap price (approximately US$ 12 kg^−1^) ^1, 6^, the chemically-synthesized vanillin is not suitable for food and beverage industry. Hence, the price of “natural” vanillin is almost 250 times higher than the synthetic vanillin ^7^.

Microbial synthesized vanillin from natural substrates is classified as “natural” vanillin under European and US food legislation ^8–9^. For instance, ferulic acid and eugenol are commonly realized as substrates for the biocatalytic production of vanillin ^10–15^. In comparison, *de novo* biosynthesis of vanillin from simple sugars such as glucose is a more attractive alternative because sugars are less expensive than ferulic acid and eugenol. Frost *et al*. harnessed the recombinant *Escherichia coli* for synthesis of vanillin from glucose via a two-step approach ^16^: vanillate was first produced by fermentation, and it was then reduced to vanillin by aryl aldehyde dehydrogenase *in vitro*. Mimicking the natural vanillin pathway from plants was also established in *E. coli* and 19.3 mg l^−1^ vanillin was achieved from glucose ^17^. However, microbial synthesis of vanillin is hindered by the host native metabolism that could rapidly convert vanillin to the byproducts such as vanillyl alcohol and vanillate. With the development of synthetic biology and metabolic engineering, researchers began to focus on addressing the instability issue of vanillin caused by the host metabolism. For example, Kunjapur *et al*. employed *E. coli* with reduced aldehyde reduction as a platform for aromatic aldehyde biosynthesis ^18^. After introducing the vanillin biosynthetic pathway, the engineered *E. coli* produced 119 mg l^−1^ vanillin from glucose ^18^. Jin-Ho Lee *et al.* identified that a single gene knockout of *NCgl0324* in *Corynebacterium glutamicum* could substantially improve the production of protocatechualdehyde and vanillin ^19^.

*Saccharomyces cerevisiae* represents the industrial “workhorse” for biochemical productions. However, *S. cerevisiae* naturally prefers alcoholic fermentation with more than 30 endogenous oxidoreductases ^20^, which is a formidable task to generate an aldehyde-accumulating yeast platform. In this study, we enabled *S. cerevisiae* to accumulate aromatic aldehydes without compromising the cell growth by systematically deleting twelve genes involved in aldehyde reduction and oxidation. Subsequently, we explored multiple coenzyme-A free vanillin biosynthetic pathways into the aldehyde-accumulating yeast for vanillin synthesis (Fig. 1). After systematic engineering to optimize the supply of NADPH and *S*-adenosylmethionine (SAM) together with metabolic reconfiguration of precursor supply, the best recombinant strain produced 365.55 ± 7.42 mg l^−1^ under shake-flasks, which represents the highest titer from glucose achieved in any microbes. Taken together, our work lays a foundation for the future implementation of vanillin production from glucose in budding yeast. In addition, the engineered **r**educed aromatic **a**ldehyde **re**duction (RARE) yeast platform might be also applicable for synthesis of many aldehyde-related compounds (Supplementary Fig. S1).

**Figure 1.**
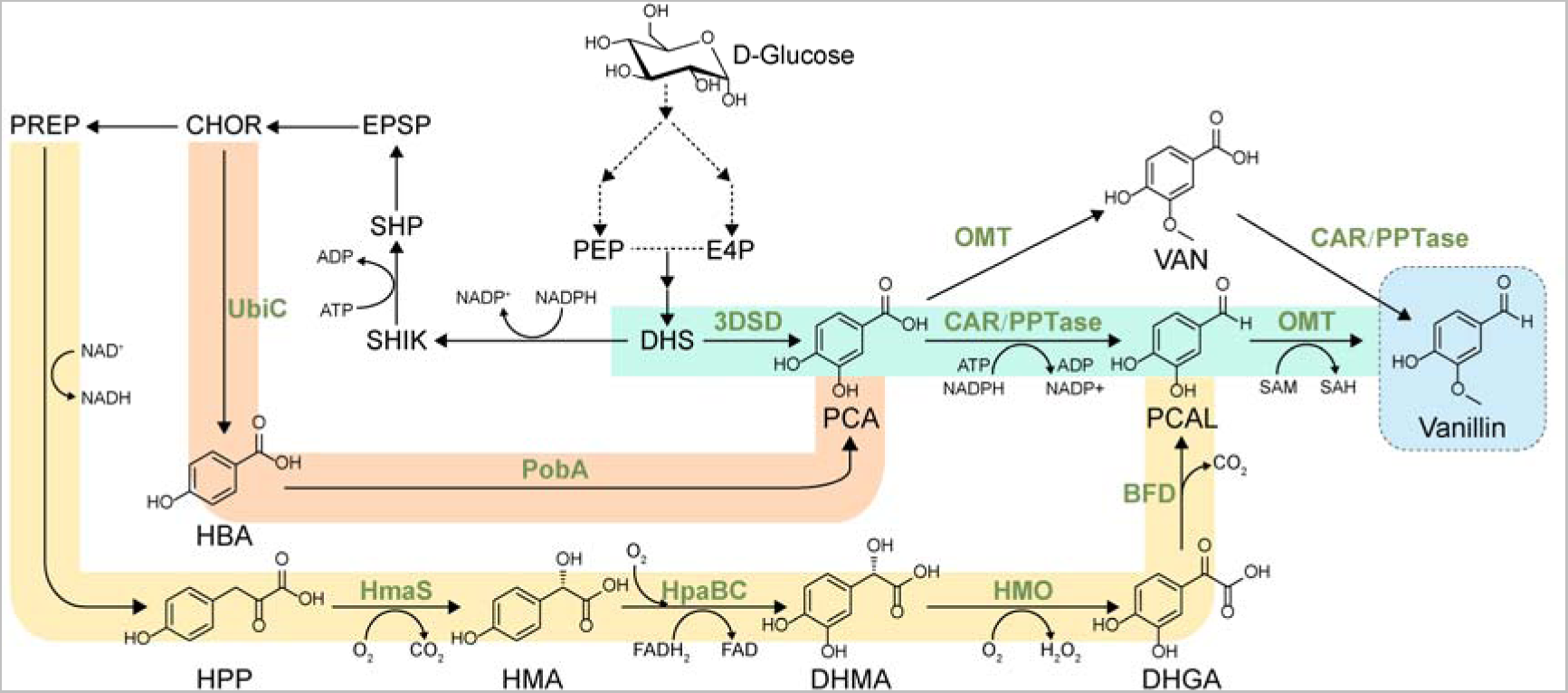
Schematic diagram of *de novo* synthesis of vanillin in *S. cerevisiae*. Colored boxes represent different metabolic routes towards vanillin synthesis. 3DSD, 3-dehydroshikimate dehydratase; CAR, carboxylic acid reductase; PPTase, phosphopantetheine transferase, OMT, *O*-methyltransferase; UbiC, chorismate-pyruvate lyase; PobA, hydroxybenzoate hydroxylase; HmaS, hydroxymandelate synthase; HpaBC, two-component flavin-dependent monooxygenase; HMO, hydroxymandelate oxidase; BFD, benzoylformate decarboxylase. PCA, protocatechuate; PCAL, protocatechualdehyde; VAN, vanillate; HBA, hydroxybenzoic acid; HPP, hydroxyphenylpyruvate; HMA, hydroxymandelate; DHMA, 3,4-dihydroxymandelate; DHGA, 3,4-dihydroxyphenylglyoxylate; PEP, phosphoenolpyruvate; E4P, D-erythrose 4-phosphate; DHS, 3-dehydroshikimate; SHIK, shikimate; SHP, shikimate-5-phosphate; CHOR, chorismate; EPSP, 5-enolpyruvylshikimate-3-phosphate; PREP, prephenate.

## Results

### Development of a yeast platform for vanillin accumulation

*De novo* synthesis of vanillin from glucose in *S. cerevisiae* was first reported by Evolva, and 45 mg l^−1^ vanillin was obtained ^21^. It was found that the majority of the vanillin product was converted to vanillyl alcohol due to the endogenous oxidoreductase activities in budding yeast ^21^. Subsequent screening of single knockout of 29 known or hypothesized oxidoreductases revealed that *Adh6* represented the most crucial gene for vanillin reduction, and a 50%-decreased ability of vanillin reduction was achieved by Δ*adh6* ^21^. More recently, our group addressed the reduction of retinal to retinol in budding yeast by a combined deletion of four alcohol dehydrogenases (ADHs) ^22^, indicating that multiple gene deletion is necessary to improve aldehyde accumulation in yeast ^23^. In this study, we implemented the same set of gene deletions (*adh6*, *adh7*, *sfa1* and *gre2*, Fig. 2a) together with the replacement of *hfd1* with *ubiC* from *E. coli* to prevent the vanillin oxidation and a dramatic improvement in the vanillin accumulation was observed (Fig. 2b).

**Figure 2.**
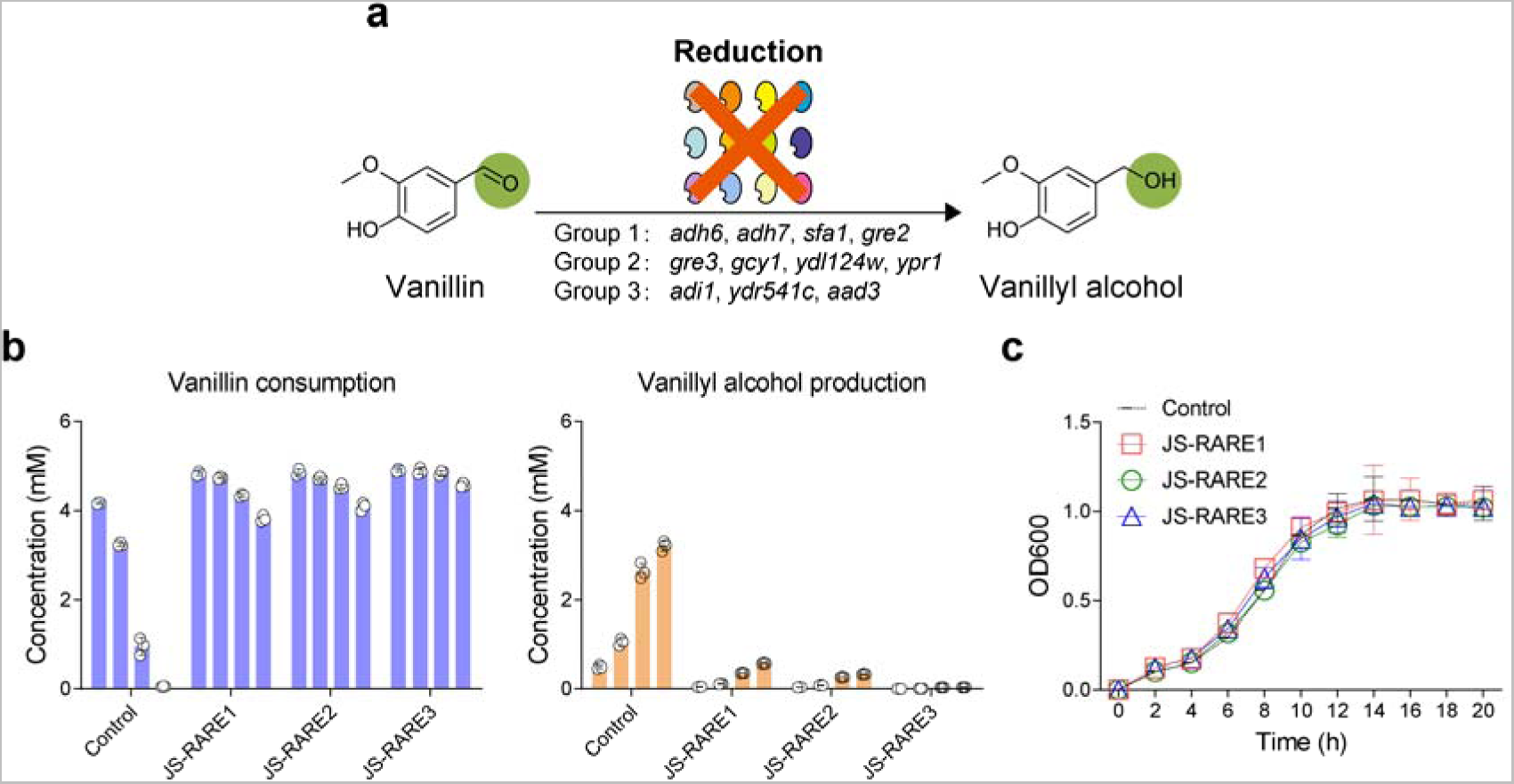
Design and build a reduced aromatic aldehyde reduction (RARE) yeast platform. **a**, Schematic diagram showing different sets of oxidoreductases for deletion to reduce vanillin conversion to vanillyl alcohol. **b**, The vanillin stability test using engineered *S. cerevisiae.* Cells were harvested after 24 h cultivation in SC media. Equal amounts of cells were resuspended into potassium-phosphate buffer (pH 8.0) with 2% glucose + 5 mM of vanillin to a final OD600 of 10. Samples were monitored at regular intervals (4, 8, 24, 48 h) using gas chromatography. **c**, The deletion of oxidoreductases did not compromise the growth of engineered yeasts. Control, the parental strain of JS-CR (2M).

Besides ADHs, deletions of additional aldo-keto reductases (AKRs) and aldehyde reductases (ALDRs) in *E. coli* are required to further prevent the aromatic aldehyde reduction ^18^. Thus, we proceeded to delete a second set of four genes (*gre3*, *gcy1*, *ydl124w*, and *ypr1*) belongs to AKR family ^24^ (Fig. 2a). The third set of three genes (*ari1*, *ydr541c*, and *aad3*) related to ALDR family ^23^ were chosen for deletion (Fig. 2a). The deletion events were confirmed by diagnostic PCR in strain JS-RARE3 (Supplementary Fig. S2). As shown in Fig. 2b, strain JS-RARE3 had the best performance to prevent vanillin reduction, and vanillyl alcohol was barely detectable by gas-chromatography even after 48 h. In addition, we found that our engineered yeasts could also substantially improve the accumulation of hydroxybenzaldehyde and protocatechualdehyde (Supplementary Fig. S3). Despite the extensive genetic engineering steps, we found that all the engineered strains showed similar growth profiles to the parental strain (Fig. 2c), suggesting that these oxidoreductases might be functionally-redundant and dispensable under the normal cultivation. To the best of our knowledge, our work represented one of the pioneering studies to investigate simultaneous deletion of more than 10 oxidoreductases for rendering the yeast with improved aldehyde-accumulating ability, and the engineered yeast had a clear advantage over the previously established yeast platform by single gene knockout ^21^.

### The RARE yeast platform enables the accumulation of *de novo* synthesized vanillin

Upon the construction of the RARE yeast platform, we next proceeded to investigate whether *de novo* synthesized vanillin could also be accumulated without forming alcohol byproduct. As shown in Fig. 1, we examined the well-established artificial vanillin biosynthetic pathway that contains 3-dehydroshikimate dehydratase (3DSD) from *Podospora anserina* (encoded by *AroZ* gene) ^25^, *O*-methyltransferase (OMT) from *Homo sapiens*, and carboxylic acid reductase (CAR) from *Segniliparus rotundus* together with *Nocardia* phosphopantetheine transferase (PPTase) ^26^. In brief, 3DSD first converts 3-dehydroshikimate (DHS) into protocatechuate. Depending on the relative enzyme kinetics and availability of cofactors, protocatechuate can be converted into protocatechualdehyde or vanillate. The final step in the pathway is either the conversion of protocatechualdehyde to vanillin by OMT or the vanillate reduction by CAR.

We first constructed two plasmids, namely, pRS423-AroZ/OMT and pRS425-CAR/PPTase. Based on the plasmid results, the main products produced by the control strain were protocatechuic alcohol and vanillyl alcohol, and only a trace amount of vanillin was accumulated (Fig. 3a). In contrast, JS-RARE3 predominantly produced vanillin (79.35 ± 0.39 mg l^−1^) with no detectable amount of vanillyl alcohol (Fig. 3a), confirming that *de novo* synthesized vanillin is relatively stable in our RARE yeast. Subsequently, we decided to integrate the vanillin biosynthetic modules into the yeast chromosomes for stable genetic inheritance. In particular, AroZ-OMT and CAR-PPTase were integrated at the sites of *rox1* and *bts1*, respectively. As shown in Fig. 3b, chromosomal integration of AroZ-OMT and CAR-PPTase substantially improved the vanillin titer, reaching 104.07 ± 3.74 mg l^−1^ vanillin after 120 h cultivation. However, we observed a substantial accumulation of protocatechuate (Fig. 3b), indicating that the CAR-mediated reduction of protocatechuate might be rate-limiting. As CAR might possibly be subjected to a substrate inhibition, we therefore further examined 3DSD from *Bacillus cereu*s (encoded by *AsbF* gene) with a lower activity in yeast. However, the reduced flux of protocatechuate synthesis did not improve vanillin synthesis, and the strain with AsbF from *B. cererus* only resulted in 77.17 ± 7.24 mg l^−1^ vanillin after 120 h (Fig. 3b).

**Figure 3.**
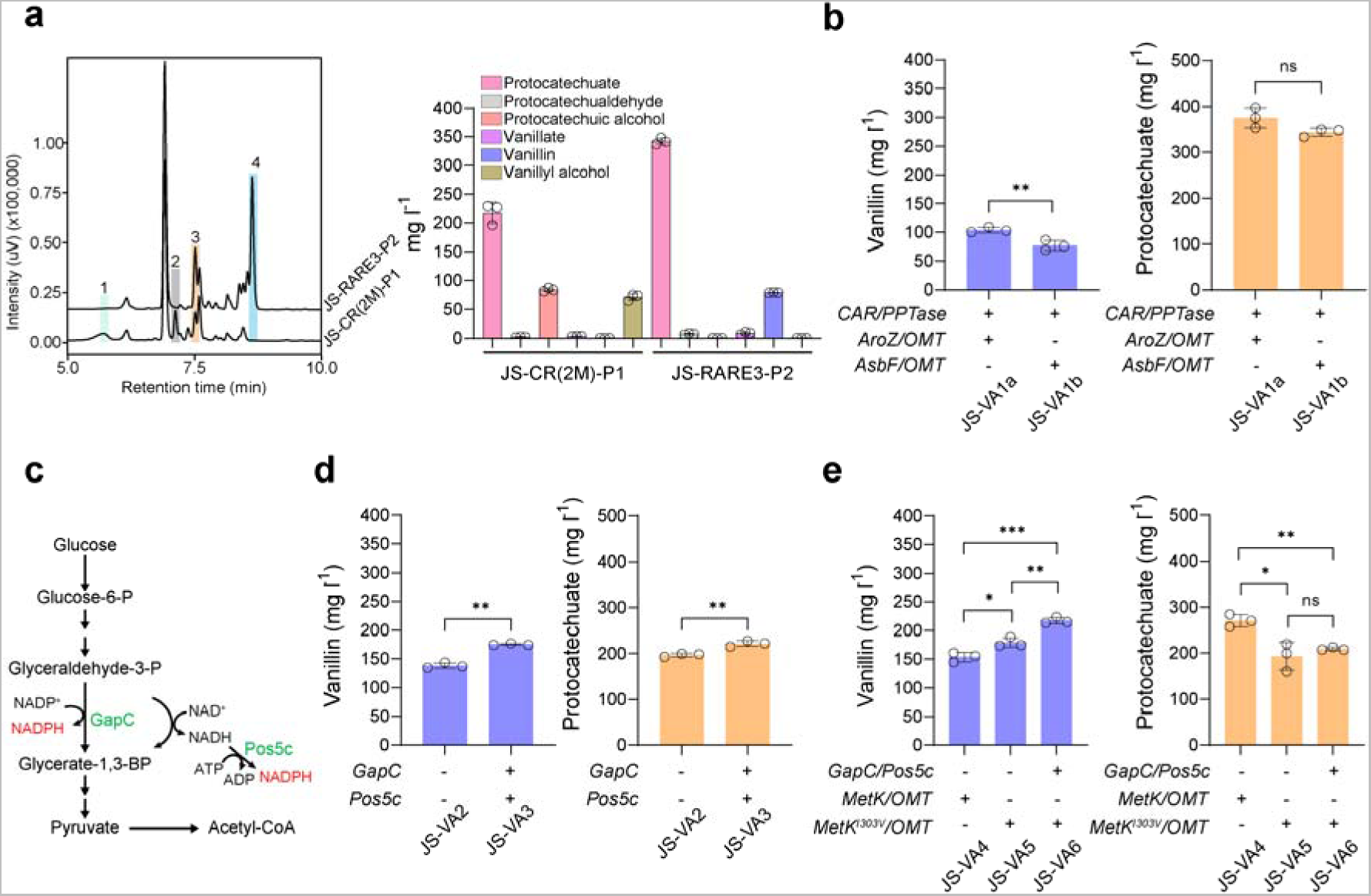
*De novo* synthesis of vanillin using 3DSD-mediated pathway in *S. cerevisiae*. **a**, Representative HPLC results showing vanillin levels in different strains and product distribution profile. Plasmid pRS423-AroZ/OMT and pRS425-CAR/PPTase were transformed into strain JS-CR(2M) and JS-RARE3, respectively. Peak 1, protocatechuic alcohol; peak 2, vanillyl alcohol; peak 3, protocatechualdehyde; peak 4, vanillin. **b**, The vanillin and protocatechuate levels in strains with chromosomal 3DSD-mediated pathway. 3DSD encoded by *AroZ* from *P. anserina* and *AsbF* from *B. cereu*s. **c**, Schematic illustration of different strategies in improving the NADPH supply. **d**, The vanillin and protocatechuate levels in strains with engineered NADPH metabolism. **e**, The vanillin and protocatechuate levels in strains with engineered SAM cycle. Cells were grown in SC medium with 2% glucose, and samples were measured after 120 h of cultivation. All experiments were performed in triplicate and the data represent the mean value with standard deviation. Statistical analysis was carried out by using two-tailed unpaired Student’s t-test (\**P* < 0.05, \*\**P* < 0.01, \*\*\**P* < 0.001).

To further improve the CAR activity for vanillin synthesis, we attempted to integrate an additional copy of PPTase from difference sources (*Sfp* from *Bacillus subtilis* and *EntD* from *E. coli*) ^21^ at the *ypl062w* site of JS-VA1a, the resulting strain JS-VA2 increased the vanillin titer to 138.50 ± 3.81 mg l^−1^ (Supplementary Fig. S4). However, the total amount of intermediates such as protocatechuate, vanillate and protocatechualdehyde was substantially reduced (Supplementary Fig. S4), indicating that disruption of *ypl062w* might affect the overall productivity of 3DSD-mediated pathway.

### Systematic engineering of the cofactor supply for improving vanillin synthesis

Considering the 3DSD-mediated vanillin biosynthetic pathway is limited because the CAR activity is not optimal in yeast, we next proceeded to optimize the abundance of cofactors (NADPH and ATP) for improving the CAR step. As shown in Fig. 3c, NADPH-dependent glyceraldehyde-3-phosphate dehydrogenase (GapC) from *Clostridium acetobutylicum* ^27^ and a cytosol-relocalized NADH kinase (Pos5c) ^28^ from *S. cerevisiae* were introduced to improve the NADPH supply, and the resulting strain was designated as JS-VA3. Upon overexpressing GapC and Pos5c, strain JS-VA3 produced 175.29 ± 1.16 mg l^−1^ vanillin and 221.66 ± 5.12 mg l^−1^ protocatechuate (Fig. 3d). In addition, the intermediate vanillate was reduced from 20.01 ± 1.26 mg l^−1^ to 9.32 ± 0.92 mg l^−1^ (Supplementary Fig. S5). Although overexpressing GapC and Pos5c could increase the vanillin production, it still did not completely solve the problem of protocatechuate accumulation. Considering CAR from *Mycobacterium abscessus* was recently reported to have a relatively high activity for the vanillate reduction ^29^, this alternative version of CAR might potentially solve the bottleneck of 3DSD-mediated vanillin production.

Next, we further proceeded to engineering the SAM supply cycle to improve the vanillin synthesis. The accumulation of protocatechuate instead of vanillate suggested that the activity of OMT from *H. sapiens* for protocatechuate methylation was clearly insufficient. The *E. coli* SAM synthetase (MetK with a mutation of I303V) was previously reported to have a 4-fold increase in activity with decreased product inhibition ^30^. By introducing the genes *MetK* or *MetK^I^*^303*V*^ together with an additional copy of *OMT* into the *yjl064w* site of strain JS-VA2, we found that the resulting strain JS-VA4 and JS-VA5 produced 153.74 ± 6.75 mg l^−1^ and 179.04 ± 6.82 mg l^−1^ vanillin, respectively (Fig. 3e). Further combined with GapC-Pos5c into JS-VA5 resulted in additional 21.36% improvement of vanillin in JS-VA6 (Fig. 3e), achieving 217.29 ± 4.65 mg l^−1^ vanillin after 120 h cultivation. The vanillate was substantially reduced in JS-VA6 over that in JS-VA5 (Supplementary Fig. S5), whereas no significant difference of protocatechuate was observed in JS-VA6 (Fig. 3e). Taken together, these results proved that the activation of SAM cycle by overexpressing MetK^I303V^ can substantially promote the methylation process mediated by OMT. However, further engineering the SAM cycle to improve the SAM supply is still required to address the vanillin productivity at the OMT-mediated methylation step ^31^.

### Implementation of a dual synthetic pathway to further enhance vanillin production

Recently, our group have demonstrated that two component flavin-dependent monooxygenase (HpaBC) from *E. coli*, hydroxymandelate synthase (HmaS) from *Amycolatopsis orientalis*, hydroxymandelate oxidase (HMO) from *Streptomyces coelicolor* and benzoylformate decarboxylase (BFD) from *Pseudomonas putida* allowed the synthesis of protocatechualdehyde ^32^. To further harness the metabolic flux from L-tyrosine synthesis, we transplanted this coenzyme A independent pathway into yeast for vanillin synthesis (Fig. 4a). As the HpaBC pair containing HpaB from *Pseudomonas aeruginosa* and HpaC from *Salmonella enterica* had a better activity when heterologously expressed in *S. cerevisiae* ^31^, we therefore replaced HpaBC from *E. coli* with PaHpaB-SeHpaC. Subsequently, we integrated HmaS-OMT at the *pha2* locus to restrict L-phenylalanine synthesis, PaHpaB-SeHpaC at the *are1* locus to minimize ergosteryl ester levels ^33^, and BFD-HMO at the *gdh1* locus to further reduce NADPH consumption ^34^.

**Figure 4.**
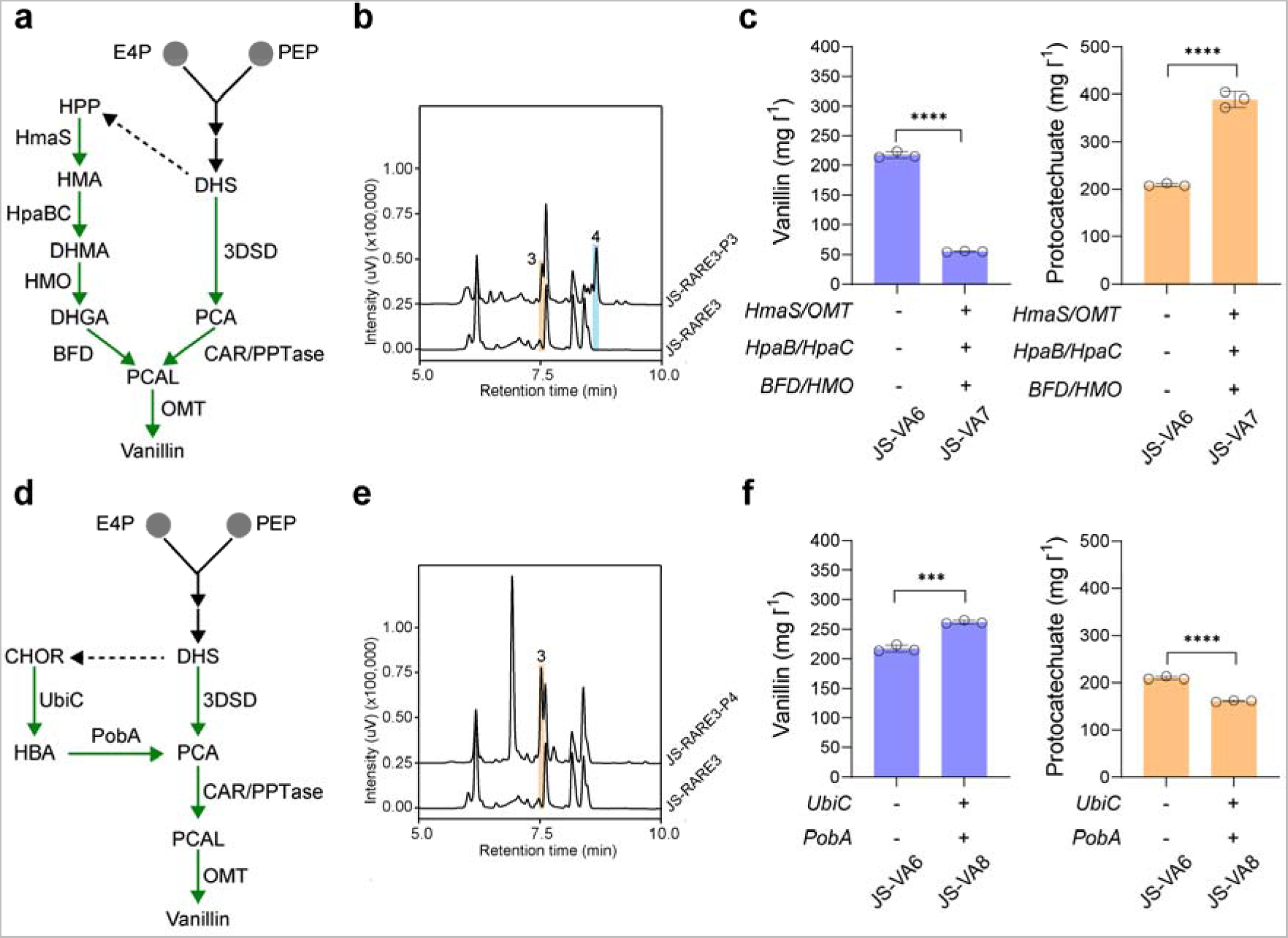
Dual synthetic pathway for enhanced vanillin synthesis in yeast. **a**, Schematic of HmaS-mediated pathway for synthesizing vanillin. **b**, Representative HPLC results of HmaS-mediated pathway for vanillin synthesis. Strains harboring plasmids with HmaS-OMT-HpaBC-HMO-BFD were used. **c**, The vanillin and protocatechuate levels in strains harboring dual 3DSD and HmaS-mediated vanillin pathway. **d**, Schematic of UbiC-PobA pathway for synthesizing vanillin. **e**, Representative HPLC results of UbiC-PobA coupled with CAR-PPTase for protocatechualdehyde synthesis. Strains harboring plasmids with UbiC-PobA-CAR-PPTase were used. **f**, The vanillin and protocatechuate levels in strains harboring dual 3DSD and UbiC-PobA vanillin pathway. Cells were grown in SC medium with 2% glucose, and samples were measured after 120 h of cultivation. All experiments were performed in triplicate and the data represent the mean value with standard deviation. Statistical analysis was carried out by using two-tailed unpaired Student’s *t*-test (\*\*\**P* < 0.001, \*\*\*\**P* < 0.0001).

We first examined plasmid-based expression of HmaS-OMT-HpaBC-HMO-BFD for vanillin production. It was found that HmaS-mediated synthetic vanillin pathway functioned in yeast based on the HPLC results (Fig. 4b), reaching up to 19.81 ± 1.54 mg l^−1^ vanillin under shake-flask cultivation (Supplementary Fig. S6). However, the engineered strain with dual 3DSD and HmaS-mediated vanillin pathway (strain JS-VA7) only produce 55.19 ± 0.60 mg l^−1^ vanillin, and more than 1.8-fold of protocatechuate was accumulated over that of the parental strain JS-VA6 (Fig. 4c). These findings indicated that the intermediates from HmaS-mediated pathway might inhibit the CAR activity. When strain JS-VA6 was supplemented with an additional 1 mM hydroxymandelate, the vanillin level was reduced from 53.31 ± 1.54 mg l^−1^ to 26.41 ± 1.40 mg l^−1^ in small-scale shake tubes for 48h (Supplementary Fig. S7), suggesting that the CAR activity might be competitively inhibited by hydroxymandelate analogs. Overall, HmaS-mediated biosynthetic pathway was functional but not compatible with the CAR-mediated vanillin pathway.

Alternatively, we also investigated chorismate pyruvate-lyase (UbiC from *E. coli*) ^35^ coupled with hydroxybenzoate hydroxylase (PobA from *P. putida*) ^36^ for vanillin synthesis (Fig. 4d). Based on the plasmid results, protocatechualdehyde was successfully produced upon introducing UbiC-PobA-CAR-PPTase (Fig. 4e). As shown in Fig. 4f, further integration of UbiC-PobA at the *gdh1* locus resulted in 262.27 ± 2.36 mg l^−1^ vanillin in strain JS-VA8, which represents 20.70% improvement over that of strain JS-VA6. Surprisingly, the protocatechuate level in JS-VA8 was substantially reduced over that in JS-VA6 (Fig. 4f), whereas there was no significant difference in the vanillate level (Supplementary Fig. S8). Since the UbiC-PobA module is a source module for additional protocatechuate synthesis, it is likely that metabolic rebalancing between 3DSD and UbiC-PobA pathways occurred as otherwise it would have elevated levels of both protocatechuate and vanillin.

### Metabolic reconfiguration for further enhanced vanillin production

To further enhance the performance of vanillin production, we also attempted to reconfigure the yeast central metabolism for an improved precursor supply of D-erythrose 4-phosphate (E4P). Recently, a phosphoketalose-based pathway (Xfpk-Pta) by providing more E4P was proven to be effective in increasing aromatic chemical productions (Fig. 5a) ^37–38^. In addition, acetyl-CoA from Xfpk-Pta pathway could be used for ATP generation, which also favorably drives the CAR step. As shown in Fig. 5b, the vanillin titer in strain JS-VA9 was further increased by 34.32% upon introducing a phosphoketolase from *Bifidobacterium breve* (BbXfpk) and a phosphotransacetylase from *Clostridium kluyveri* (CkPta), reaching 352.28 ± 7.03 mg l^−1^ (17.61 ± 0.35 mg per g glucose) under shake flasks.

**Figure 5.**
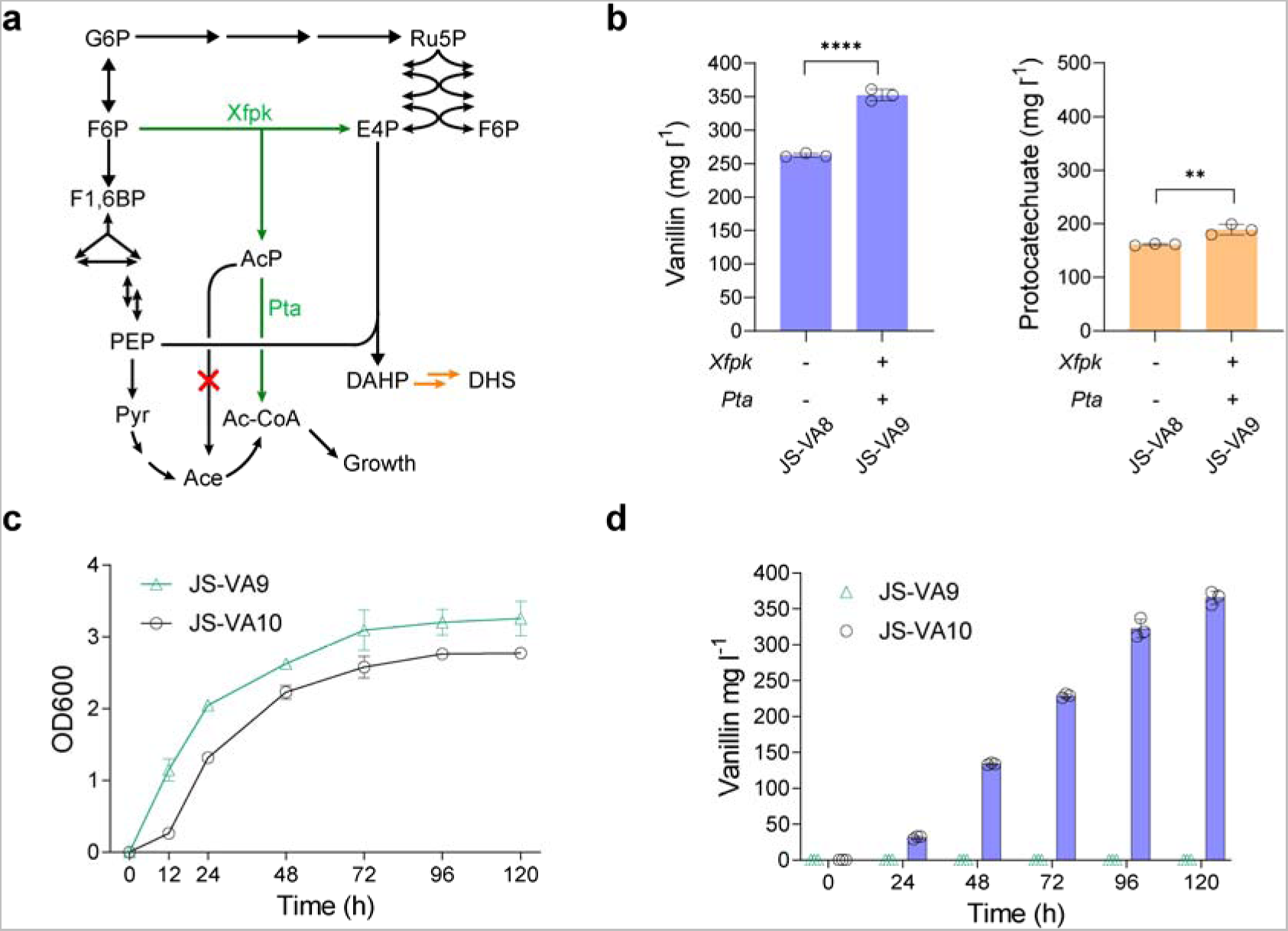
Metabolic reconfiguration for further enhanced vanillin production. **a**, Schematic overview of Xfpk-Pta pathway for improving the precursor supply of E4P. **b**, The effect of Xfpk-Pta pathway on the vanillin and protocatechuate productions. The glycerol-1-phosphatase GPP1 was deleted to minimize acetate formation (marked with a red cross). Cells were grown in SC medium with 2% glucose, and samples were measured after 120 h of cultivation. **c**, The growth curve of strain JS-VA9 and JS-VA10 in shake-flasks. Strain JS-VA10 was a derivative of JS-VA9 with Ubi-K15N degron fused to the N-terminal of Gal80. Both strains were cultivated in YPD medium containing 20LJg l^−1^ glucose. **d**, Vanillin produced by strain JS-VA9 and JS-VA10 in YPD media. All experiments were performed in triplicate and the data represent the mean value with standard deviation. Statistical analysis was carried out by using two-tailed unpaired Student’s t-test (\*\**P* < 0.01, \*\*\*\**P* < 0.0001).

To evaluate the performance of the engineered strain in yeast-peptone-dextrose (YPD) medium, we further engineered the copper-inducible system with Ubi-K15N tagged Gal80 ^39^, to reduce the half-life of Gal80. As shown in Fig. 5c, the OD600 of JS-VA10 was lower than that of the control strain JS-VA9 (no vanillin production in YPD medium). Strain JS-VA10 resulted in 365.55 ± 7.42 mg l^−1^ after 120 h cultivation (Fig. 5d). Based on our toxicity assay as shown in Supplementary Fig. S9, the yeast cells can tolerate <1 g l^−1^ vanillin and multiple genetic modifications of oxidoreductase deletions did not change the yeast tolerance to vanillin. However, the titer of vanillin in YPD was only slightly improved over that in SC medium, indicating that *de novo* synthesized vanillin might be even more toxic than extracellularly supplemented vanillin. Therefore, the bottleneck for industrial production of vanillin would be the toxicity of vanillin to the cells ^21^ and future industrial-scale vanillin production in yeast would require strain engineering to improve the yeast tolerance to vanillin.

## Discussion

Aromatic aldehydes are used widely as flavors and fragrances, and serve as intermediates for alkaloid synthesis ^40, 41^. However, the main barrier to microbial synthesis of aromatic aldehydes is the instability of aldehydes, which can be rapidly converted into corresponding alcohols or acids. In this study, we have constructed a RARE yeast platform that accumulates an industrially-relevant aromatic aldehyde of vanillin. The optimal aldehyde-accumulating result was achieved by combined deletion of 11 genes, spanning the ADH, AKR, and ALDR super-families. Despite all these deletions, the growth rate of the engineered strains remained nearly uncompromised. Besides, we also observed that a variety of aromatic aldehydes (hydroxybenzaldehyde, protocatechualdehyde) could also be accumulated in the engineered yeast, suggesting that the RARE yeast platform might be applicable for future synthesis of many aldehyde-derived compounds.

Natural vanillin biosynthetic pathways typically reply on ferulic acid as the precursor, which is converted to vanillin by a single hydratase/lyase type enzyme designated as vanillin synthase (VpVAN) ^42^, or by a CoA-dependent pathway comprising feruloyl-CoA synthetase (FCS) and enoyl-CoA hydratase/aldolase (ECH) ^17^. In budding yeast, the endogenous FDC1 and PAD1 could rapidly decarboxylate phenylacrylic acids such as ferulic acid, caffeic acid and coumaric acid ^43^. In this study, we used artificial vanillin biosynthetic pathways to bypass the intermediate of ferulic acid. In particular, we first assembled a synthetic pathway comprising 3DSD, OMT, CAR, and PPTase. We found that *AroZ* gene from *P. anserina* was more effective for protocatechuate production than *AsbF* gene from *B. cereus*. However, the primary limiting factor for 3DSD-mediated vanillin synthesis seems to be the insufficient activities of CAR and OMT, which resulted in a substantial amount of protocatechuate accumulation. Nevertheless, the byproduct of protocatechuate is also a high-value chemical with potential therapeutic applications ^44–46^.

We also investigated the hydroxymandelate degradation pathway (HmaS, HMO and BFD) ^47–48^ and UbiC-PobA for improving the vanillin production. It was found that the heterologous hydroxymandelate degradation pathway was functional in budding yeast, but this pathway was not compatible with 3DSD-mediated vanillin synthesis as the CAR activity would be repressed by hydroxymandelate analogs. Noteworthily, UbiC-PobA was proven to an effective strategy to further improve the vanillin titer in the strain equipped with 3DSD-mediated vanillin pathway and the synthetic yeast factory with dual vanillin biosynthetic pathway produced 262.27 ± 2.35 mg l^−1^ in shake-flasks. Further introducing Xfpk from *B. breve* and Pta from *C. kluyveri* to improve the precursor E4P supply enabled 352.28 ± 7.03 mg l^−1^ vanillin in shake flasks, which represents the highest vanillin titer from glucose.

According to the literature, nonglucosylated vanillin beyond the 0.5-to 1-g l^−1^ scale would severely hamper the growth of *S. cerevisiae* ^21^. The remaining bottleneck for vanillin overproduction in yeast is the intrinsic toxicity of vanillin to the host cells. Adaptive laboratory evolution (ALE) has proven a useful strategy to acquire desired phenotypes with accumulation of beneficial mutations under selective pressure. Moreover, global transcription machinery engineering (gTME) by mutagenesis of the transcription factor could lead to dominant mutations that confer increased tolerance and more efficient glucose conversion to ethanol ^49^. In the future, ALE and gTME to reprogram the global cell metabolism might be implemented to improve the vanillin tolerance in budding yeast. In addition, diploid *S. cerevisiae* or other yeast species with a better tolerance might be used as the starting chassis for industrial-scale vanillin production. Alternatively, *in situ* vanillin recovery using resin with high selectivity and loading capacity might be used to address the toxicity issue of vanillin. Therefore, we believe that sustainable production of natural vanillin from glucose would be eventually achieved by continuous efforts in metabolic engineering, synthetic biology and process optimization.

## Method

### Strains, culture media and reagents

*S. cerevisiae* BY4741 derived JS-CR(2M) ^50^ was used as the initial chassis for constructing all the subsequent strains. The YPD medium (10 g l^−1^ yeast extract, 20 g l^−1^ peptone and 20 g l^−1^ glucose) was used for normal cultivation of yeast cells, and synthetic complete (SC) medium with appropriate dropouts was used for yeast cells with different auxotrophic markers. *E. coli* DH5α was used as the recipient strain for cloning plasmids, and the strains carrying the plasmid were cultured at 37°C in Luria-Bertani broth with 100 μg ml^−1^ ampicillin. All restriction enzymes, T4 ligase, Taq polymerase and High-fidelity Phusion polymerase were obtained from New England Biolabs (Beverly, MA, USA). Gel extraction kit and plasmid purification kit were purchased from BioFlux (Shanghai, China). Antibiotics, 5-fluoroorotic acid and oligonucleotides were purchased from Sangon Biotech (Shanghai, China). The standard vanillin (Cat. No. V100115), vanillyl alcohol (Cat. No. H103777), vanillate (Cat. No. V104428), hydroxybenzaldehyde (Cat. No. H100420), hydroxybenzyl alcohol Cat. No. H107912), protocatechualdehyde (Cat. No. D108634), protocatechuic alcohol (Cat. No. D155345), and protocatechuate (Cat. No. P104382) were purchased from Aladdin Biotech (Shanghai, China). All the other chemicals were obtained from Sigma-Aldrich or otherwise stated.

### Plasmid construction

Oligonucleotides used for plasmid construction are listed in Supplementary Table S1. *HmaS* from *A. orientalis*, *OMT* from *H. sapiens*, *HpaB* from *P. aeruginosa*, *Xfpk* from *B. breve* and *Pta* from *C. kluyveri* were synthesized by GenScript (Supplementary Table S2). *MetK*, *UbiC* and *EntD* were PCR amplified from the genomic DNA of *E. coli* MG1655. *HpaC* was PCR amplified from the genomic DNA of *S. enterica* LT2. *HMO* was PCR amplified from the genomic DNA of *S. coelicolor* A3(2). *Segniliparus CAR* and *Nocardia PPTase* were kindly provided by Prof. Dunming Zhu from Tianjin Institute of Industrial Biotechnology, Chinese Academy of Sciences. *AroZ* was a gift from Prof. Eckhard Boles from Goethe University Frankfurt. *BFD* and *PobA* were PCR amplified from the genomic DNA of *P. putida* KT2440. *Sfp* and *AsbF* was PCR amplified from the genomic DNA of *B. subtilis* 168. *Pos5c* was PCR amplified from the genomic DNA of *S. cerevisiae* BY4741. *GapC* was PCR amplified from the genomic DNA of *C. acetobutylicum* ATCC 824. Plasmid pRS423-AroZ/OMT, pRS423-AsbF/OMT, pRS425-CAR/PPTase, pRS423-HpaB/HpaC, pRS425-BFD/HMO, pRS426-HmaS/OMT, pRS423-MetK/OMT, pRS423-MetK^I303V^/OMT, pRS423-Xfpk/Pta, pRS426-UbiC/PobA, and pRS425-Sfp/EntD, were all constructed via the golden-gate approach ^51^. All the plasmids used in this study are provided in Supplementary Table S3.

### Genome editing of *S. cerevisiae*

The CRISPR/Cas9 genome editing was carried out as previously described ^52^. The guide RNA (gRNA) expression plasmids were derived from an inhouse plasmid of pRS426SNR52. The standard protocol of *S. cerevisiae* transformation was carried out by electroporation with minimal modification. 50 μl of yeast cells together with approximately 2 μg mixture of genome editing cassette was electroporated in a 0.2 cm cuvette at 1.6 kV. After electroporation, cells were immediately mixed with 900 μl YPD medium and recovered a rotary shaker for 1 h. Cells were plated on SC plate with appropriate dropouts. Successful genome manipulations were confirmed by diagnostic PCR before proceeding to the next round of genetic modifications. Subsequently, gRNA expressing plasmid was eliminated via counter-selection with 1 g l^−1^ 5-fluorootic acid (5-FOA), and the Cas9-expressing plasmid was removed via a series dilution. The flowchart of yeast strain construction is provided in Supplementary Fig. S10. All the strains used in this study are provided in Supplementary Table S4.

### Shake-flask cultivation for vanillin production

For small-scale production of vanillin, experiments were carried out using 100 ml shake-flasks. 1% fresh overnight culture was inoculated into shake-flasks containing 10 ml SC medium supplemented with 20 μM copper sulfphate. The cultures were incubated at 30°C and 250 rpm for vanillin productions. For gas chromatography-flame ionization detector (GC-FID) analysis of vanillin, 100 μl of supernatant was extraction with 900 μl ethyl acetate before subjected to GC-FID analysis. 1 μl of diluted sample was injected into GC-2030 system equipped with a Rtx-5 column (30 m × 250 μm × 0.25 μm thickness). Nitrogen (ultra-purity) was used as carrier gas at a flow rate 1.0 ml min^−1^. GC oven temperature was initially held at 40°C for 2 min, increased to 45°C with a gradient of 5°C min^−1^ and held for 4 min. And then it was increased to 230°C with a gradient 15°C min^−1^.

For high-performance liquid chromatography (HPLC) analysis of vanillin, Shimadzu Prominence LC-20A system (Shimadzu, Japan) equipped with a reversed phase C18 column (150 × 4.6 mm, 2.7 μm) and a photodiode array detector was used. The samples were centrifuged and filtered through a 0.2 μm syringe filter before injected to the HPLC system. The mobile phase comprises solvent A (ddH_2_O with 0.1% trifluoroacetic acid) and solvent B (acetonitrile with 0.1% trifluoroacetic acid). The following gradient elution was used: 0 min, 95% solvent A + 5% solvent B; 8 min, 20% solvent A + 80% solvent B; 10 min, 80% solvent A + 20% solvent B; 11 min, 95% solvent A + 5% solvent B. The flow rate was set at 1 ml min^−1^. The levels of vanillin and other aromatic compounds were monitored at the absorbance of 275 nm.

### Analysis of the growth-inhibitory effect of vanillin on yeast

Fresh overnight cultures of yeast strains were inoculated into SC media supplemented with different concentrations of vanillin (0.25, 0.5, 0.75, 1.0, and 1.5 g l^−1^), whereas no additional vanillin supplementation was used as the control. The yeast cultures were then grown at 30°C on a rotary shaking incubator at 250 rpm. The OD600 was measured with regular time intervals (4, 8, 12, 16, 20, 24, and 28 h).

## Acknowledgements

This work was supported by the National Natural Science Foundation of China (grant no.: 32270087), Xiamen University (grant no.: 0660X2510200), Daan Gene (20223160A0063) and ZhenSheng Biotech.

## Author Contributions

J.Y. conceived and designed the project. Q.M. performed the experiments and collected the data. J.Y. interpreted the data. J.Y. and Q.M. wrote the manuscript.

## Supporting information

is available in the online version of the paper.

