## Supplemental information for "Systems engineering of *Saccharomyces cerevisiae* for synthesis and accumulation of vanillin"

**Supporting Figures**

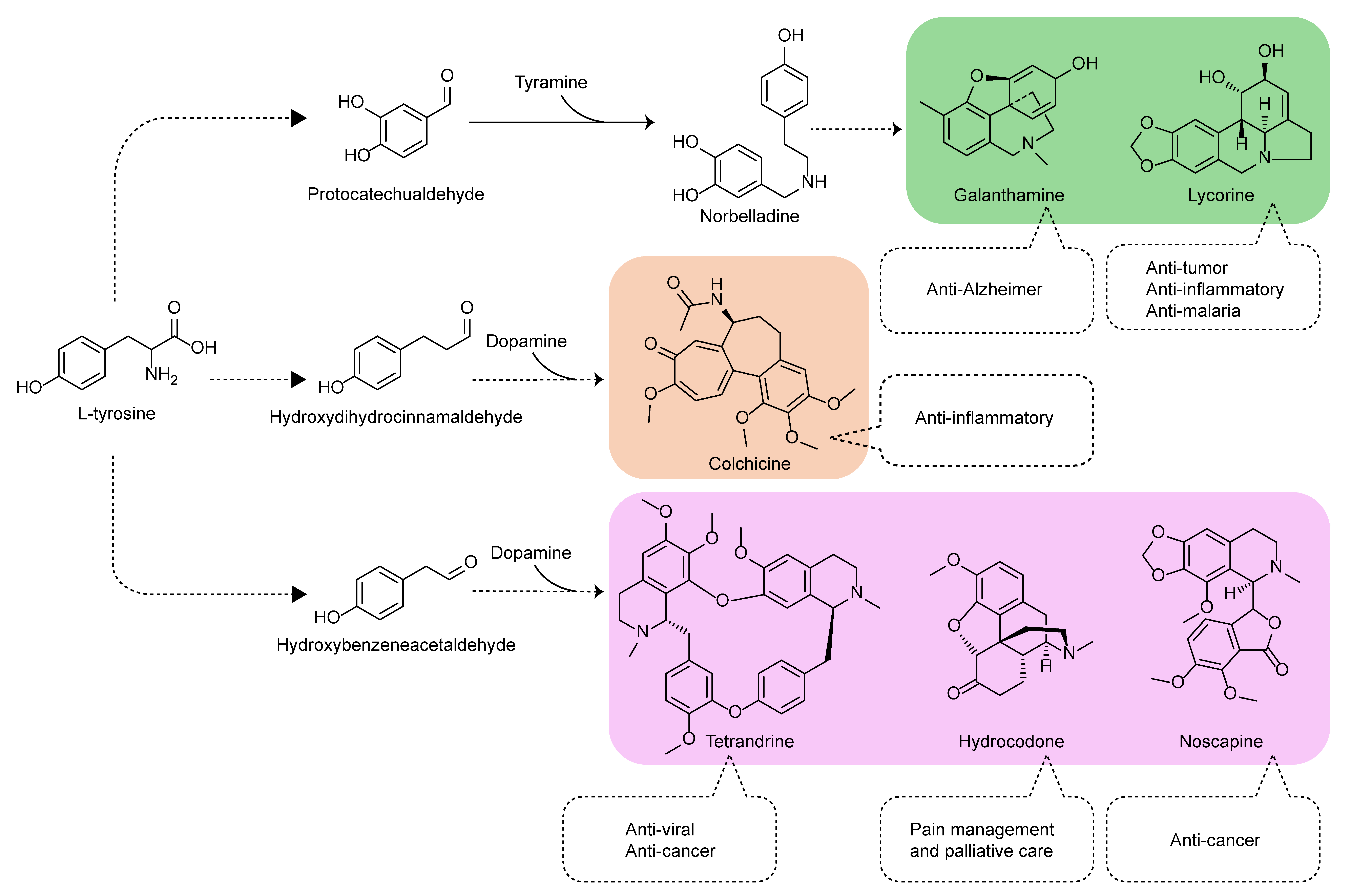

**Supplementary Fig. S1. Biosynthesis of alkaloids.**

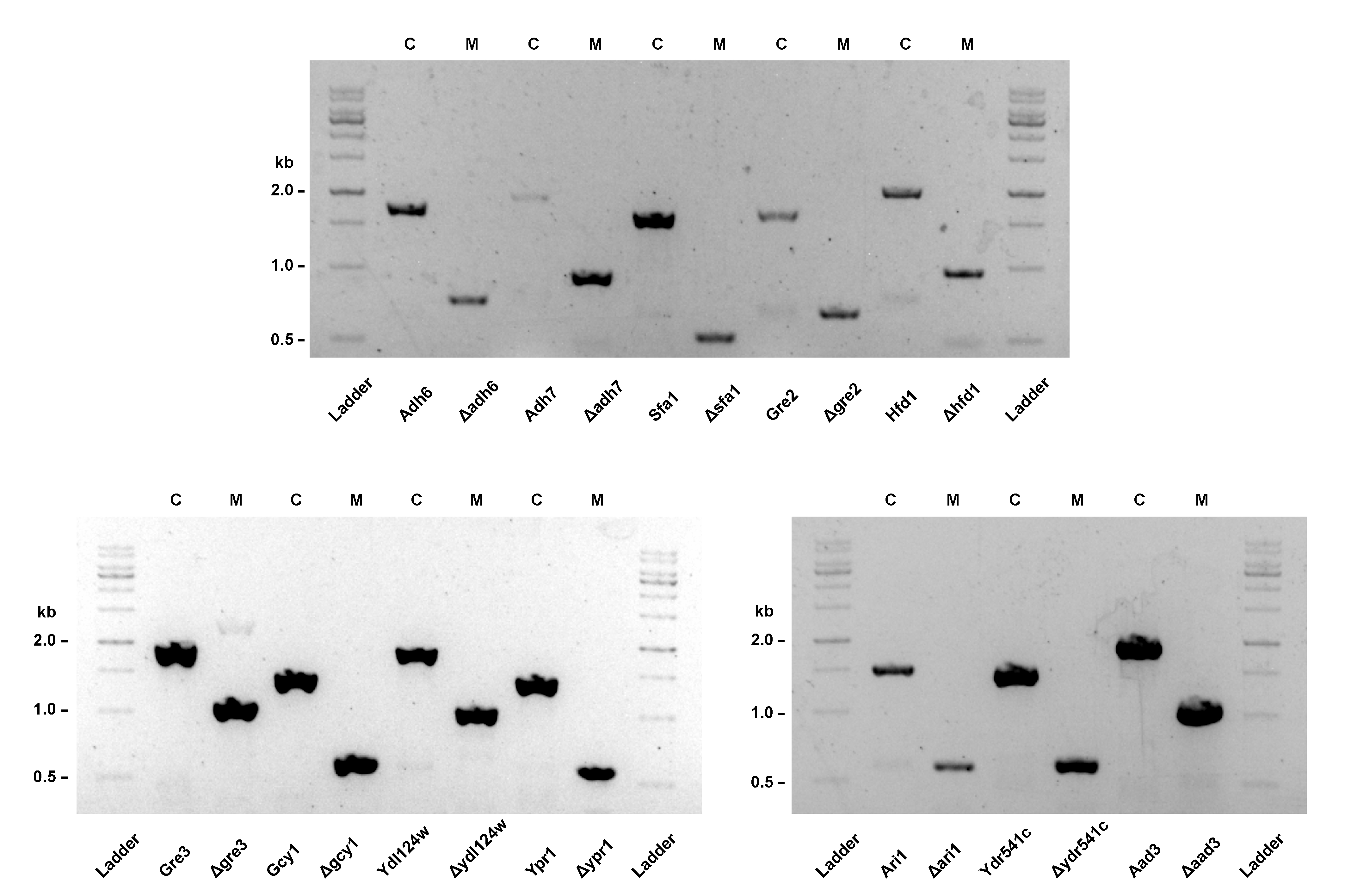

**Supplementary Fig. S2. Knockout of the endogenous oxidoreductases in budding yeast.** C, control; M, JS-RARE3.

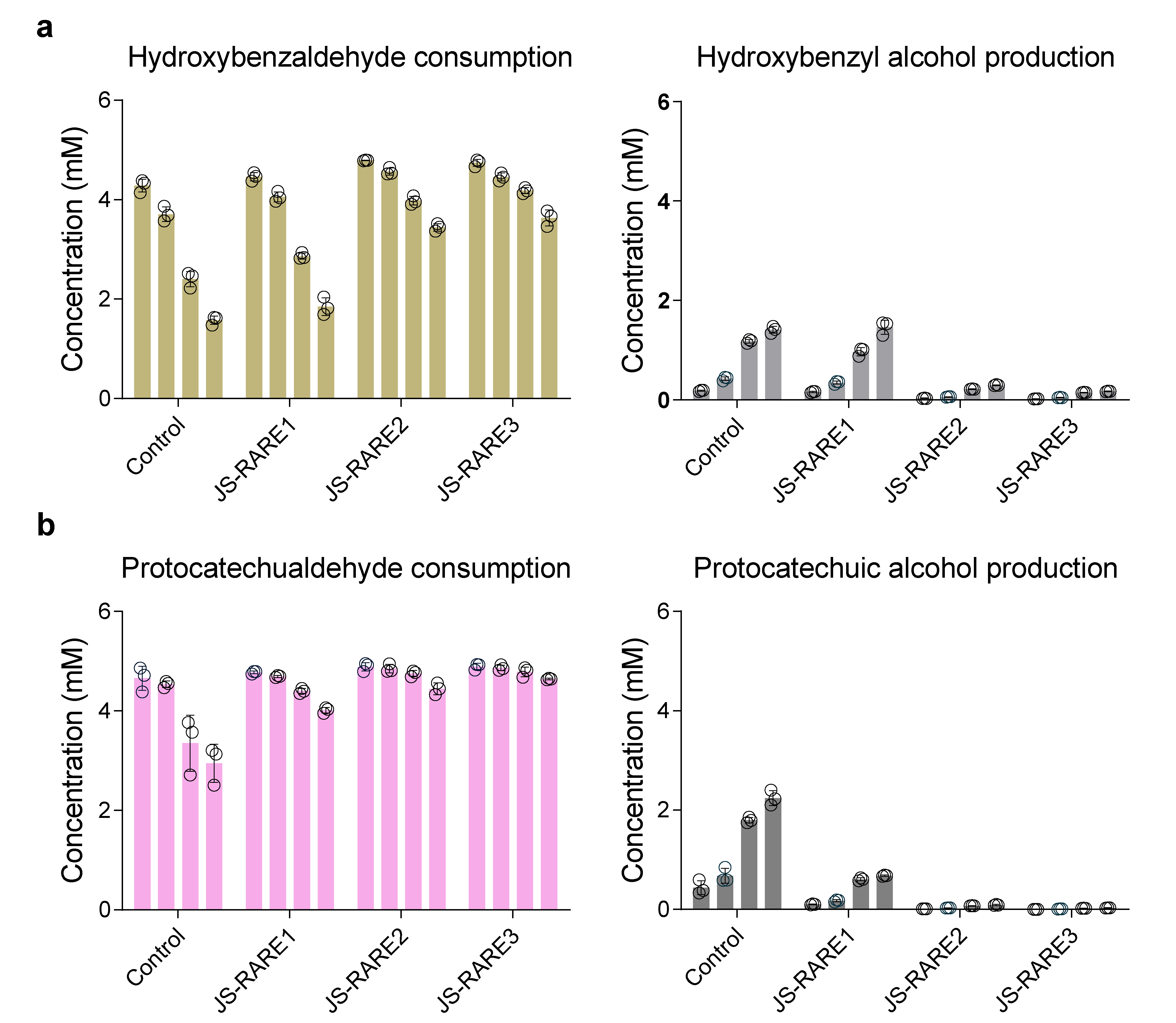

**Supplementary Fig. S3. A combination of rational gene deletions enables other aromatic aldehyde accumulation.** **a,** Hydroxybenzaldehyde stability test using the engineered *S. cerevisiae*. **b,** Protocatechualdehyde stability test using the engineered *S. cerevisiae*. Cells were harvested after 24 h cultivation in SC media. Equal amounts of cells were resuspended into KP buffer (pH 8.0) with 2% glucose + 5 mM of hydroxybenzaldehyde or protocatechualdehyde to a final OD600 of 10. Samples were periodically monitored by HPLC or GC-FID analysis for 4, 8, 24, 48 h.

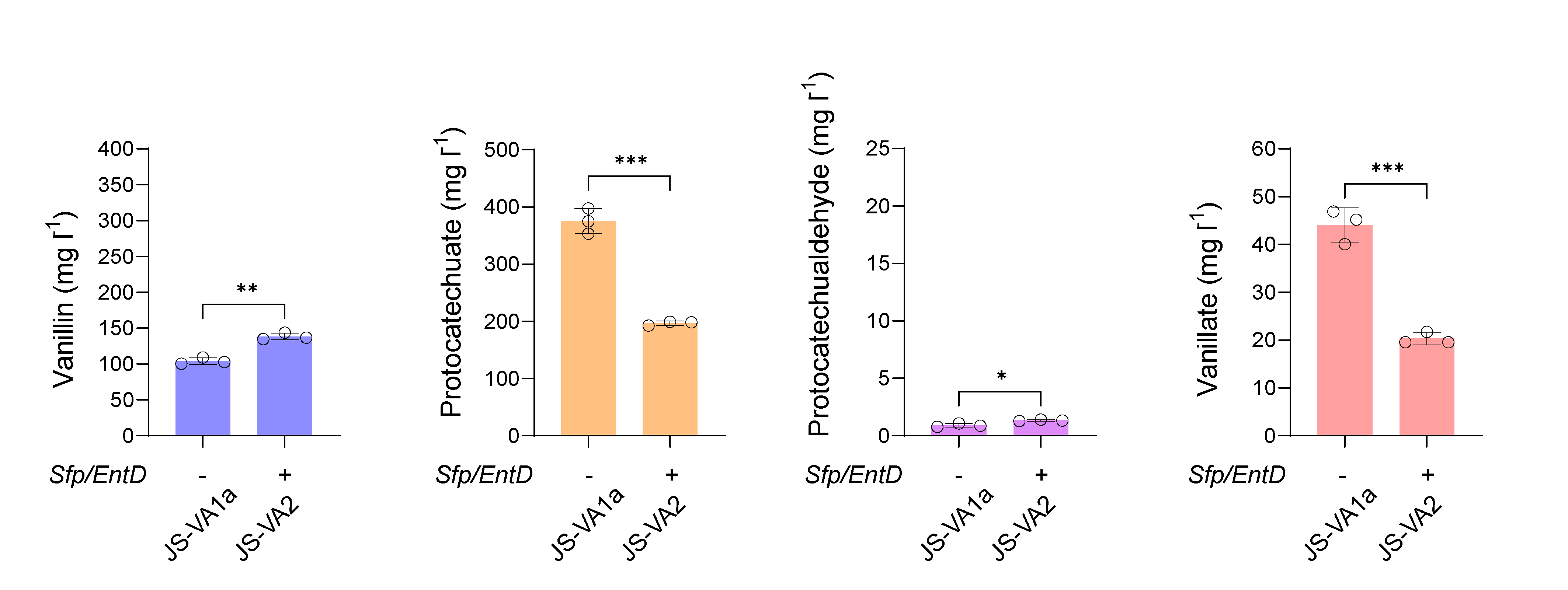

**Supplementary Fig. S4. The effect of Sfp and EntD expression on vanillin production.** The vanillin levels in strain JS-VA1a and JS-VA2 were compared. Cells were grown in SC medium with 2% glucose, and samples were measured after 120 h of cultivation. The experiments were performed in triplicate and the data represent the mean value with standard deviation.

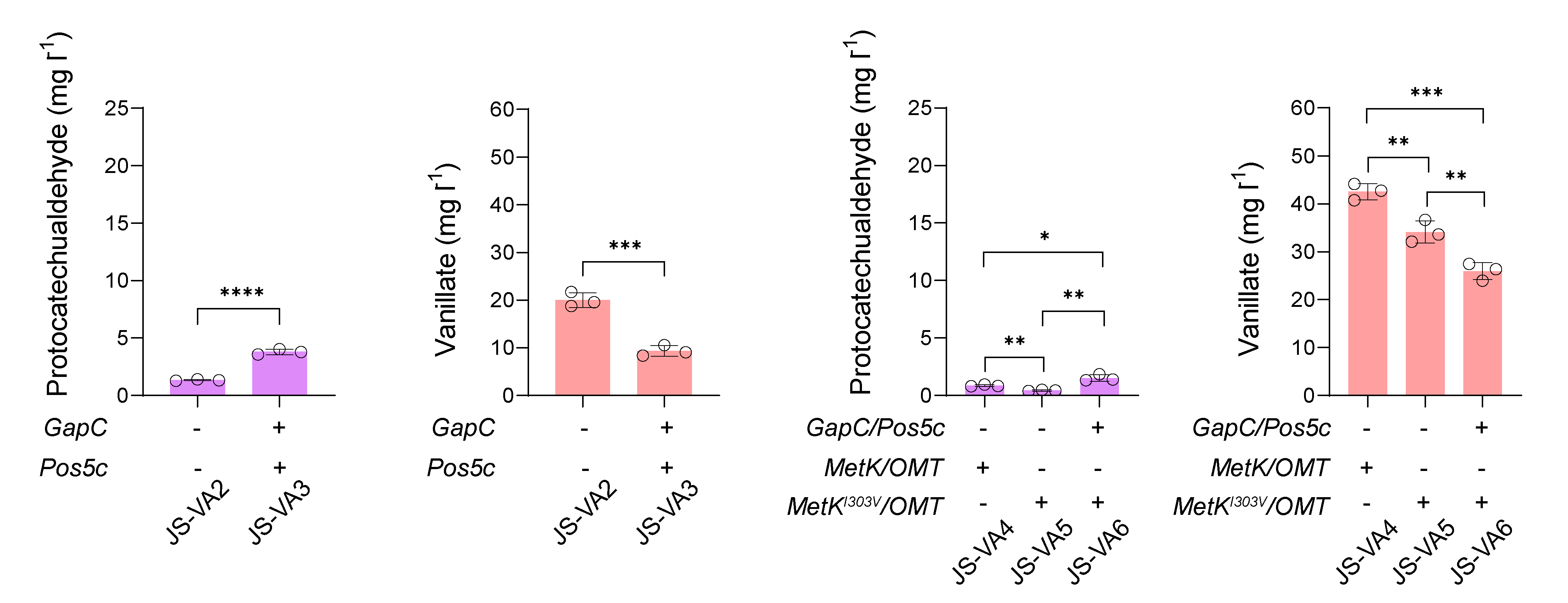

**Supplementary Fig. S5. Product profiles of protocatechualdehyde and vanillate produced by engineered yeasts of JS-VA2~6.** Cells were grown in SC medium with 2% glucose, and samples were measured after 120 h of cultivation. The experiments were performed in triplicate and the data represent the mean value with standard deviation.

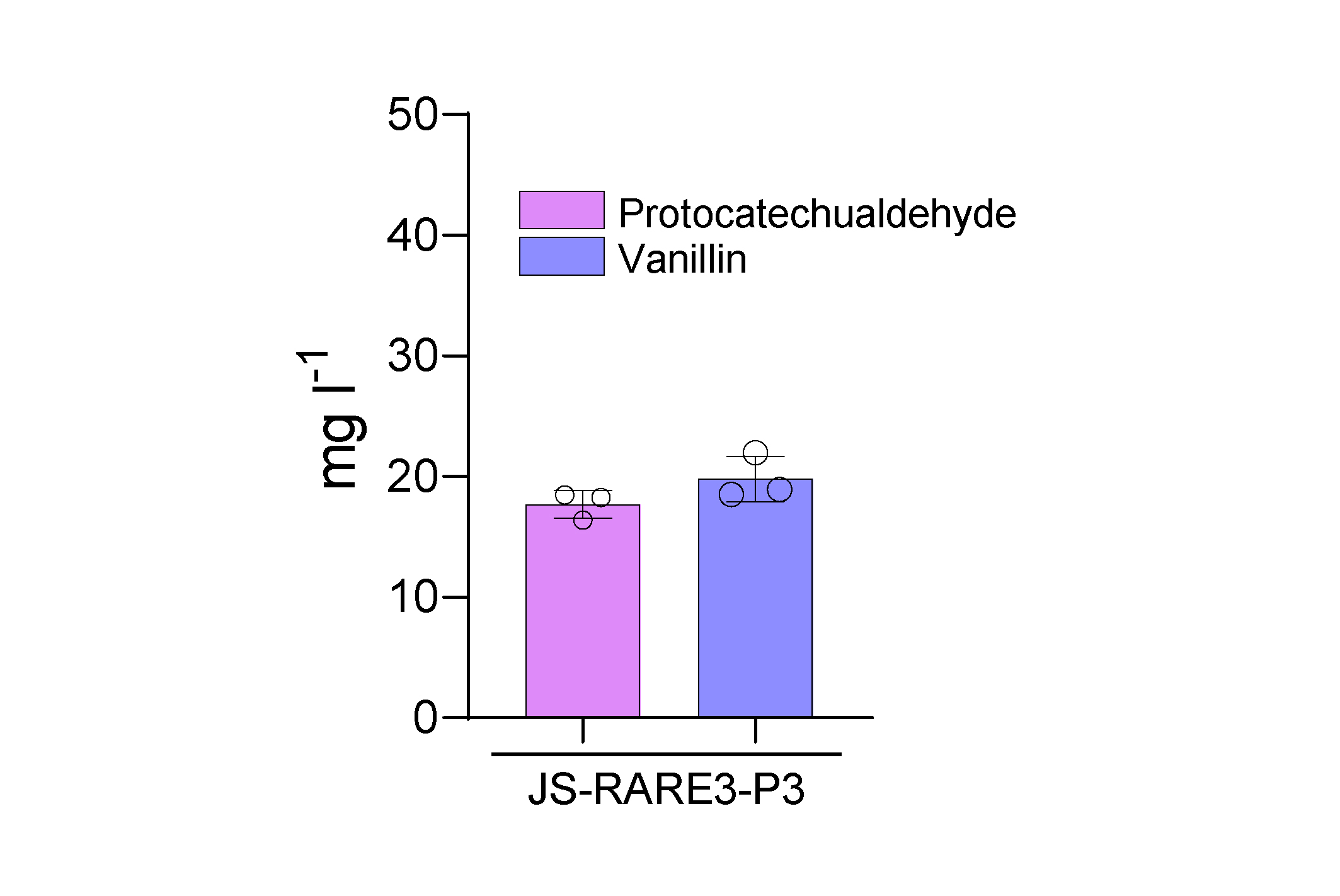

**Supplementary Fig. S6. *De novo* synthesis of vanillin from plasmid-based HmaS pathway in *S. cerevisiae*.** The strain JS-RARE3-P3 is a derivative from JS-RARE3 with plasmids pRS423-HpaB/HpaC, pRS425-BFD/HMO, and pRS426-HmaS/OMT. SC media supplemented with 2% (w/v) glucose with dropouts were used for cultivating the engineered yeasts. The experiments were performed in triplicate and the data represent the mean value with standard deviation.

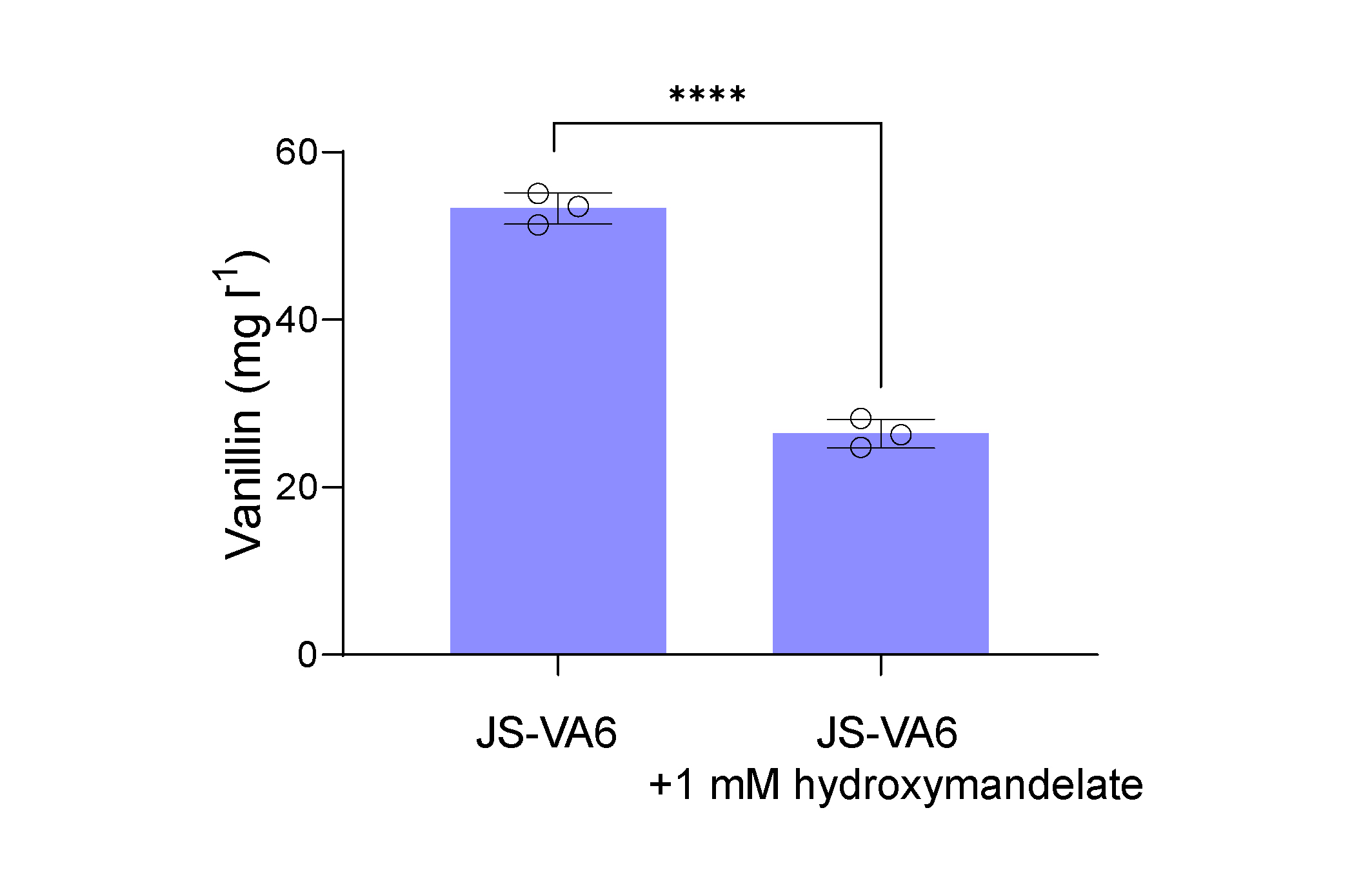

**Supplementary Fig. S7. The inhibitory effect of hydroxymandelate to the CAR-mediated vanillin biosynthetic pathway.** Comparison of the vanillin levels in *S. cerevisiae* strain (JS-VA6) with or without additional supplementation of 1 mM of hydroxymandelate. The vanillin levels in 14 ml shake tubes supplemented with 2 ml SC media were measured after 2 days. The experiments were performed in triplicate and the data represent the mean value with standard deviation.

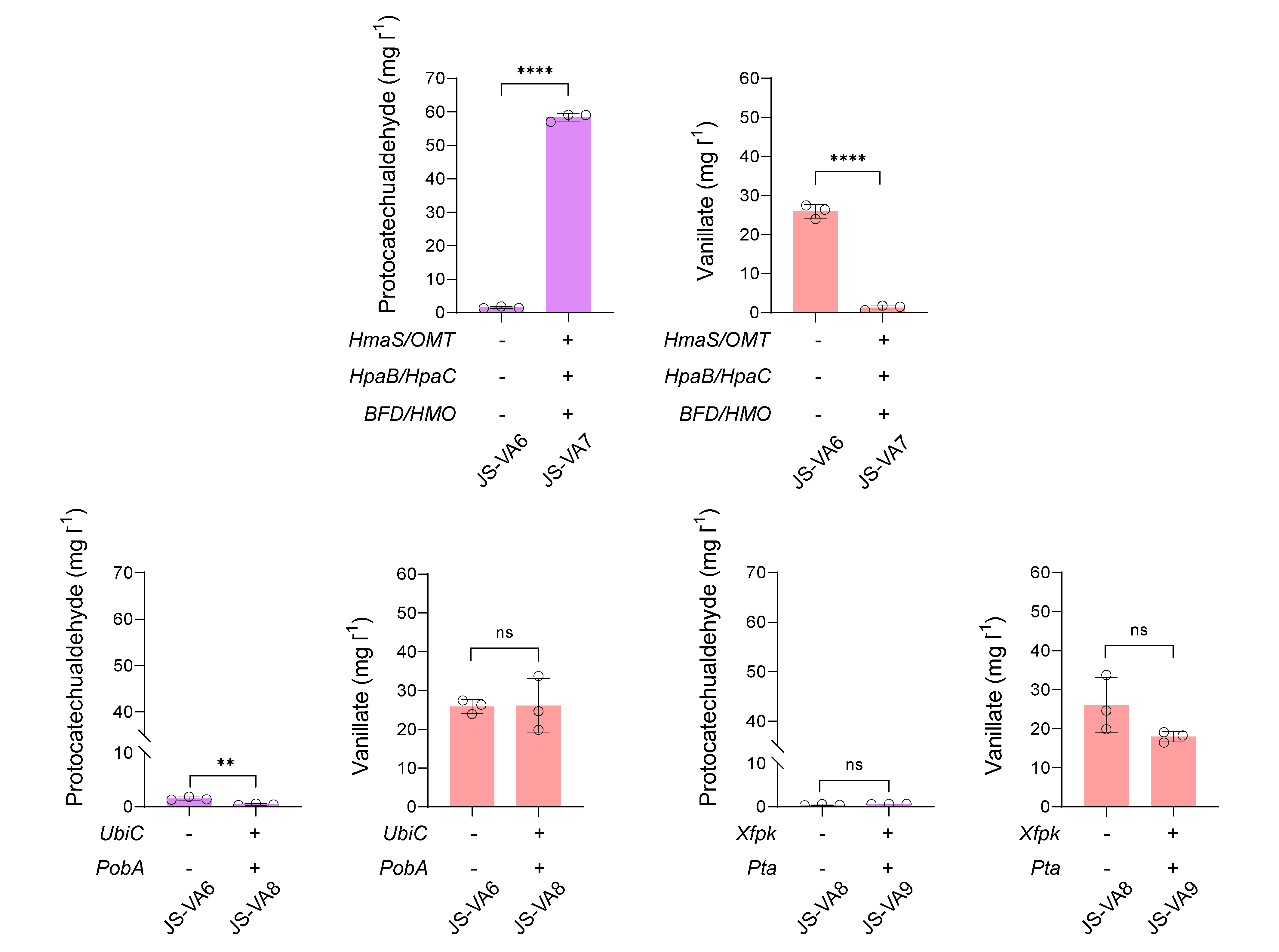

**Supplementary Fig. S8. Product profiles of protocatechualdehyde and vanillate produced by engineered yeasts of JS-VA6~9.** Cells were grown in SC medium with 2% glucose, and samples were measured after 120 h of cultivation. The experiments were performed in triplicate and the data represent the mean value with standard deviation.

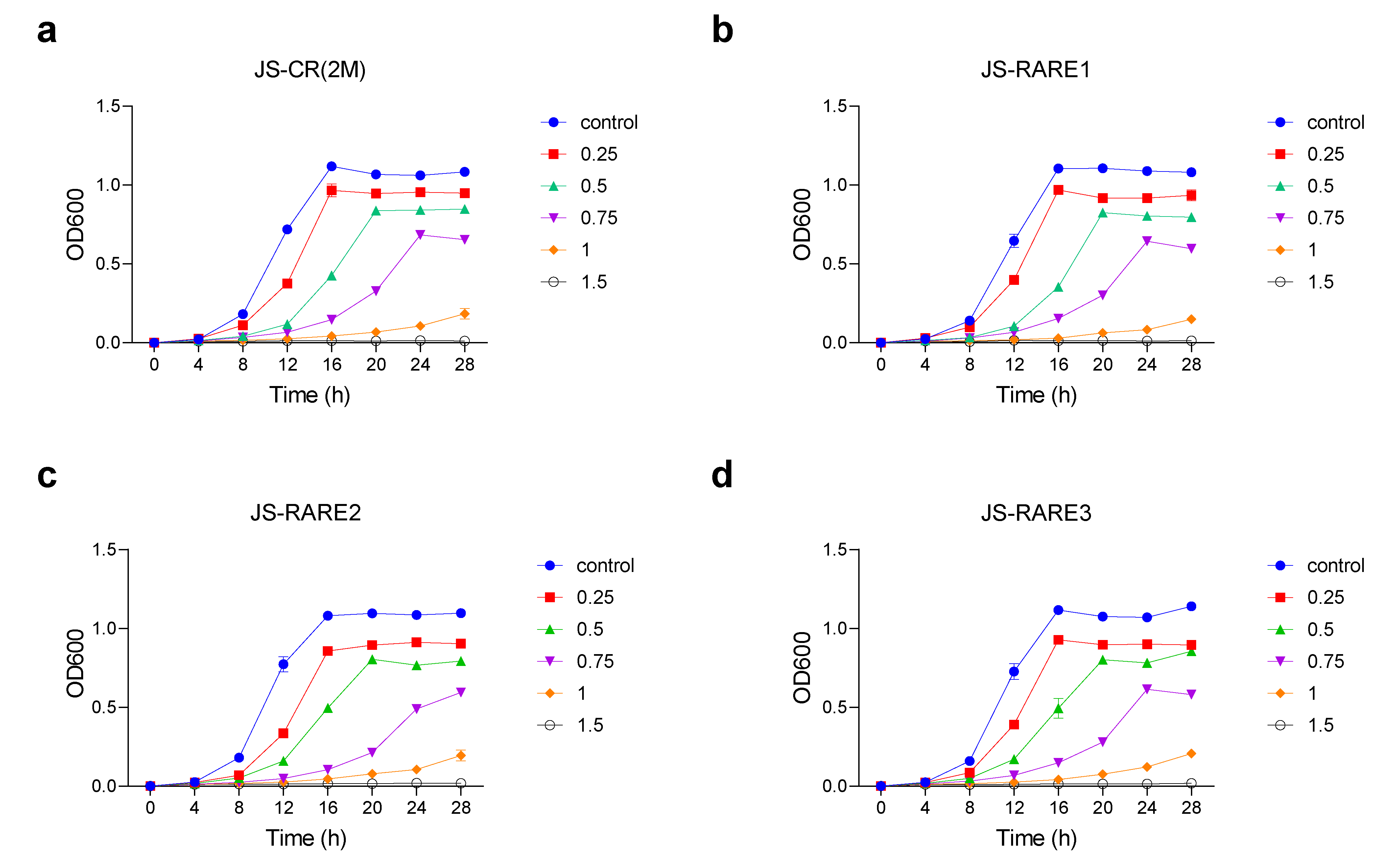

**Supplementary Fig. S9. The growth inhibitory effect of vanillin to *S. cerevisiae*.** Strain JS-CR(2M), JS-RARE1, JS-RARE2, and JS-RARE3 were treated with different concentrations of vanillin (blue dot, no vanillin supplementation; red square, 0.25 g/L vanillin; green triangle, 0.50 g/L vanillin; purple inverted triangle, 0.75 g/L vanillin; orange diamond, 1.0 g/L vanillin; black circle, 1.5 g/L vanillin).

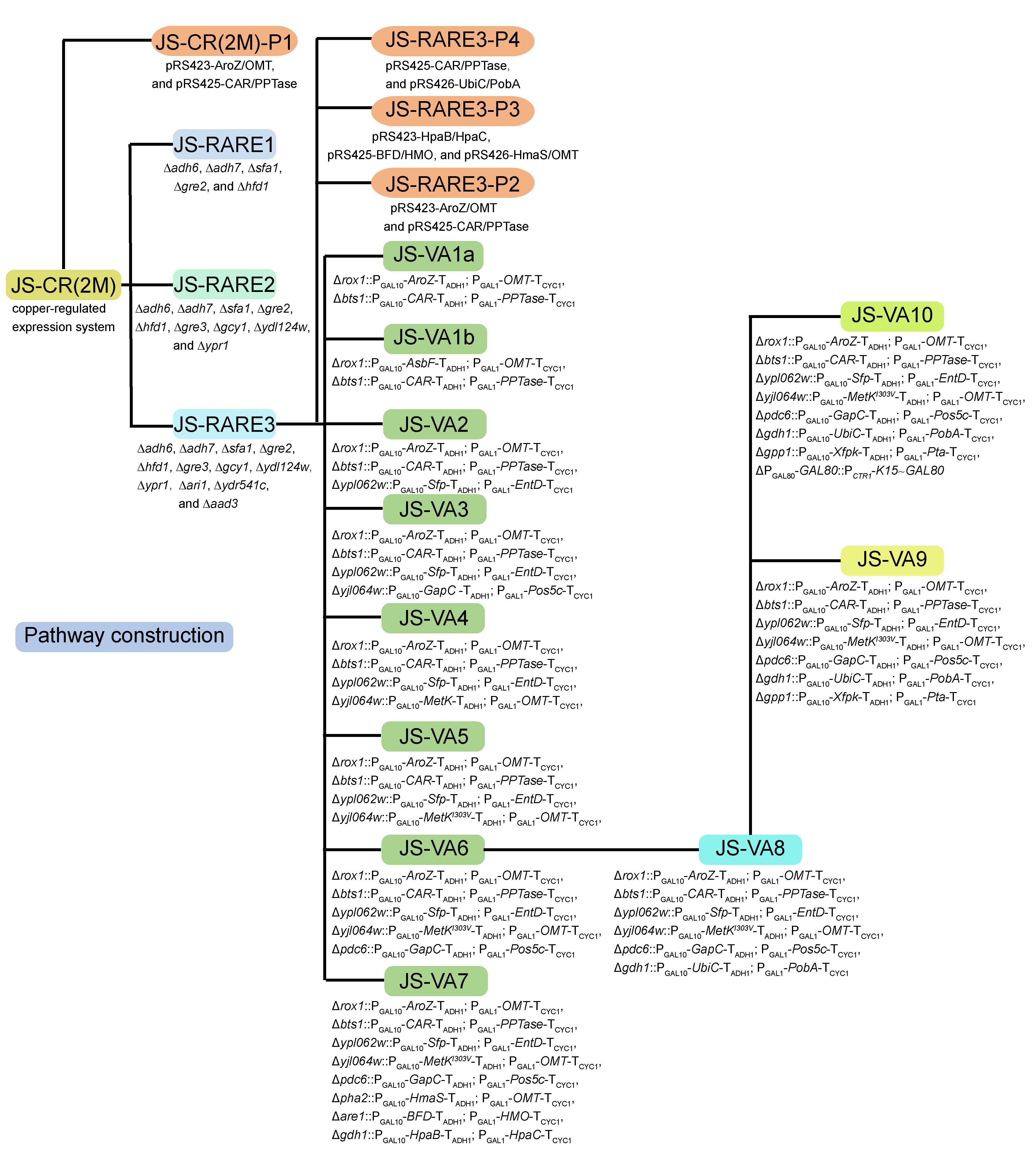

**Supplementary Fig. 10. Flowchart of yeast strain construction in this study.**

**Supporting Tables**

**Table S1.** Oligonucleotides used in this study.

| **Name** | **Description** |
| --- | --- |
| F_gRNA.gre3 | TTGGTCTCAGATGATTCTCGAACAGAGTTGGCGGTTTTAGAGCTAGAAATAG |
| F_gRNA.gcy1 | TTGGTCTCAGATGTACCGATGCTCCACTATTGAGTTTTAGAGCTAGAAATAG |
| F_gRNA.ypr1 | TTGGTCTCAGATGCAAGTGCCGAAACCCAACACGTTTTAGAGCTAGAAATAG |
| F_gRNA.ydl124w | TTGGTCTCAGATGATGTCAGATTCCCCAGCGGAGTTTTAGAGCTAGAAATAG |
| F_gRNA.ari1 | TTGGTCTCAGATGAGGAGATTTGGTGATCACGGGTTTTAGAGCTAGAAATAG |
| F_gRNA.aad3 | TTGGTCTCAGATGGACTTCTCCTAGGATTAACGGTTTTAGAGCTAGAAATAG |
| F_gRNA.ydr541c | TTGGTCTCAGATGGGTGATCAAGTCATTGACCGGTTTTAGAGCTAGAAATAG |
| F_gRNA.pdc6 | TTGGTCTCAGATGGTTGGGGTCAATCTCCTCAGGTTTTAGAGCTAGAAATAG |
| F_gRNA.pha2 | TTGGTCTCAGATGTTGAAGAGAAGGGAGAACGCGTTTTAGAGCTAGAAATAG |
| F_gRNA.are1 | TTGGTCTCAGATGCTTCACCGTAGTCTTCACCGGTTTTAGAGCTAGAAATAG |
| F_gRNA.gdh1 | TTGGTCTCAGATGACAACACCCAGAATACAGAAGTTTTAGAGCTAGAAATAG |
| F_gRNA.gpp1 | TTGGTCTCAGATGACCGTCAACATCGAATAGAGGTTTTAGAGCTAGAAATAG |
| F_gRNA.gal80N | TTGGTCTCAGATGGGACTACAACAAGAGATCTTGTTTTAGAGCTAGAAATAG |
| R_SUP4 | TTGGTCTCAAAAGAGACATAAAAAACAAAAAAAG |
| F-adh6-Del | ATGTCTTATCCTGAGAAATTTGAAGGTATCGCTATTCAATTTACCTTAG |
| R-adh6-Del | CTAGTCTGAAAATTCTTTGTCGTAGCCGACTAAGGTAAATTGAATAGCG |
| F-adh7-Del | ATGCTTTACCCAGAAAAATTTCAGGGCATCGGTATTTCCTTTACTTTGG |
| R-adh7-Del | CTATTTATGGAATTTCTTATCATAATCGACCAAAGTAAAGGAAATACCG |
| F-sfa1-Del | ATGTCCGCCGCTACTGTTGGTAAACCTATTAAGTGCATTTGCTTAAGAA |
| R-sfa1-Del | CTATTTTATTTCATCAGACTTCAAGACGGTTCTTAAGCAAATGCACTTA |
| F-gre2-Del | ATGTCAGTTTTCGTTTCAGGTGCTAACGGGTTCATTGCCACTGCCTCCC |
| R-gre2-Del | TTATATTCTGCCCTCAAATTTTAAAATTTGGGAGGCAGTGGCAATGAAC |
| F-hfd1-Del | ATATTCTAAAACCATAGCCATAGTAATTTATCACCAACATGTCACACCCCGCGTTAAC |
| R-hfd1-Del | CTTATACATCAAATAATTAATTAACCTTAAACATTACGTTTAGTACAACGGTGACGCCG |
| F-gre3-Del | ATGTCTTCACTGGTTACTCTTAATAACGGTCTGAAAATGGAGCGACCTCATGCTATAC |
| R-gre3-Del | TCAGGCAAAAGTGGGGAATTTACCATCCAACCAGGTCCAGTATAGCATGAGGTCGCTC |
| F-ydl124w-Del | ATGTCATTTCACCAACAGTTCTTTACCTTGAATAATGGATTGTACGGTA |
| R-ydl124w-Del | TTATACTTTTTGAGCAGCGTAGTTGTATTTACCGTACAATCCATTATTC |
| F-gcy1-Del | ATGCCTGCTACTTTACATGATTCTACGAAAATCCTTTCTGAGCGACCTCATGCTATAC |
| R-gcy1-Del | TTACTTGAATACTTCGAAAGGAGACCAATTTGGATGTACGTATAGCATGAGGTCGCTC |
| F-ypr1-Del | ATGCCTGCTACGTTAAAGAATTCTTCTGCTACATTAAAAGAGCGACCTCATGCTATAC |
| R-ypr1-Del | TCATTGGAAAATTGGGAAGGATCCCCACTTCATATCAACGTATAGCATGAGGTCGCTC |
| F-ari1-Del | ATGACTACTGATACCACTGTTTTCGTTTCTGGCGCAACCAGCGACCTCATGCTATAC |
| R-ari1-Del | TTAGGCTTCATTTTGAACTTCTAACATTTGCGCCGCGGTGTATAGCATGAGGTCGCT |
| F-ydr541c-Del | ATGTCTAATACAGTTCTAGTTTCTGGCGCTTCAGGTTTTAGCGACCTCATGCTATAC |
| R-ydr541c-Del | TCATAATCTGTTCTGCTTCTTCAAAATTTGGGCAGCAGTGTATAGCATGAGGTCGCT |
| F-aad3-Del | ATGATTGGGTCCGCGTCCGACTCATCTAGCAAGTTAGGAAGCGACCTCATGCTATAC |
| R-aad3-Del | CTAAACATTATTCGTACCATATTTTTGAGTCAAGGAATGTATAGCATGAGGTCGCT |
| F-bts1-Int | ATGGAGGCCAAGATAGATGAGCTGATCAATAATGATCCTGGAGCGACCTCATGCTATAC |
| R-bts1-Int | TCACAATTCGGATAAGTGGTCTATTATATATAACAATTCGCTTCGAGCGTCCCAAAACC |
| F-ypl062w-Int | ATGATAGAATTGGATTATGTAAAAGGTGAAGATACCATTGGAGCGACCTCATGCTATAC |
| R-ypl062w-Int | CTATATCGCATTCGTTGCACTCACCGTTCCCAAGAGGAGACTTCGAGCGTCCCAAAACC |
| F-rox1-Int | ATGAATCCTAAATCCTCTACACCTAAGATTCCAAGACCCAGAGCGACCTCATGCTATAC |
| R-rox1-Int | TCATTTCGGAGAAACTAGGCTAGTTTTAGCGGTGACCTCACTTCGAGCGTCCCAAAACC |
| F-yjl064w-Int | ATGACACTTGTAGTATATCTAACTCGGTTTTCTTCCACTAGAGCGACCTCATGCTATAC |
| R-yjl064w-Int | TCAGGCTAACACAATGAACAACGAGACTAGTGGTAAAGAACTTCGAGCGTCCCAAAACC |
| F-pdc6-Int | ATGTCTGAAATTACTCTTGGAAAATACTTATTTGAAAGAGAGCGACCTCATGCTATAC |
| R-pdc6-Int | TTATTGTTTGGCATTTGTAGCGGCAGTCAATTGCGCTTGCTTCGAGCGTCCCAAAACC |
| F-pha2-Int | ATGGCCAGCAAGACTTTGAGGGTTCTTTTTCTGGGTCCCGAGCGACCTCATGCTATAC |
| R-pha2-Int | TTATTTGTGATAATATCTCTCATTTCTGGGGAATGTACCCTTCGAGCGTCCCAAAACC |
| F-are1-Int | ATGACGGAGACTAAGGATTTGTTGCAAGACGAAGAGTTTGAGCGACCTCATGCTATAC |
| R-are1-Int | TCATAAGGTCAGGTACAACGTCATAATGATACTGGGCCCCTTCGAGCGTCCCAAAACC |
| F-gdh1-Int | ATGTCAGAGCCAGAATTTCAACAAGCTTACGAAGAAGTTGAGCGACCTCATGCTATAC |
| R-gdh1-Int | TTAAAATACATCACCTTGGTCAAACATAGCATCAGAGACCTTCGAGCGTCCCAAAACC |
| F-gpp1-Int | ATGCCTTTGACCACAAAACCTTTATCTTTGAAAATCAACGAGCGACCTCATGCTATAC |
| R-gpp1-Int | TTACCATTTCAACAAGTCATCCTTAGCGTATAAGTAGTCCTTCGAGCGTCCCAAAACC |
| Ubi-K15N_int_fwd | TAGAAAATAAAAAAAAGTGTATTATATTTGACATTCAAAATGCAGATTTTCGTCAAGAC |
| Ubi-K15N_int_rev | CCGACTCTTATGGGAGCTGCATTAGGCACGGTTGAGACACCAGAACCCTTAACCAAAG |
| F-adh6v | ACAGCCACTCTCGTCACGGC |
| R-adh6v | CCGATACCCTATGAACGTGC |
| F-adh7v | GATACGTTTGGCTCTGTTGC |
| R-adh7v | CACTGTTGTCGAGAGATTC |
| F-sfa1v | AGACATGCGGTGTGTGGGTC |
| R-sfa1v | GTTAGGAACAGGCGAGGTC |
| F-gre2v | ACATTGTTGTACGCTATAG |
| R-gre2v | GCTTATCTGAACGTTTCTC |
| F-hfd1v | CTTAGAGGAAATGGAACAAC |
| R-hfd1v | GAAAGGTTACTTATACATC |
| F-gre3v | CGCAGATACTGTAAATGCCG |
| R-gre3v | ATGGCTAGTGCTATCATTGC |
| F-ydl124wv | CTGTAACGATTCGCACCATATC |
| R-ydl124wv | GCGAGTTTTGAGGACGATTC |
| F-gcy1v | ATGCCTGCTACTTTACATG |
| R-gcy1v | CTTCTGGTGGCCTATCTTTG |
| F-ypr1v | CCATGCCTGCTACGTTAAAG |
| R-ypr1v | ATGAAGGAGAAGAAGATTCG |
| F-ari1v | ACCTTAGACCTTGCAACATC |
| R-ari1v | TTAGGCTTCATTTTGAACTTC |
| F-ydr541cv | CAGCAACAATGGCCAGTCGC |
| R-ydr541cv | TCATAATCTGTTCTGCTTCTTC |
| F-aad3v | CAATATCAGCGAAGAAGAGG |
| R-aad3v | CTCTACCACCTTCGTCAGAG |
| M1_bts1_F300 | CGCCATCTCTACTCACTCC |
| M1_bts1_R300 | TGTCTAGGGATGGCTCACG |
| M2_ypl062w_F300 | CACACCTAATGATCTGATGC |
| M2_ypl062w_R300 | AGTGTGATCTTGACTGGTTC |
| M3_rox1_F300 | CTCGTATTGTCTTGCCGGTG |
| M3_rox1_R300 | TATCGATAATATAGGTATAC |
| M4_yjl064w_F200 | CGTATAAAGAAACTATGTTG |
| M4_yjl064w_R200 | CTATGTCATGGTTCCCAGAC |
| Pdc6_F800 | CTTTCCTCACGGACCATACG |
| Pdc6_R0 | GTTTGGCATTTGTAGCGGC |
| Pha2_F200 | CGTACTACATCATCTGCGAC |
| Pha2_R200 | CGGTGCGGCCCCGCCACCAC |
| Are1_F200 | CTGGCTTGGCCATCAAATACC |
| Are1_R200 | GTCACCTGCAAACTCTTCTTG |
| Gdh1_F200 | GTATACGTAATCTAAGTAAG |
| Gdh1_R200 | GTCCAATCGATGCTTACATAC |
| Gpp1_F100 | TCTTTCGTAAGTATCTCTTG |
| Gpp1_R150 | GAAAAGGATGCATTACATCG |
| AroZ_P1_fwd | TTGGTCTCATGAAAACAATGCCAAGTAAACTGGCTAT |
| AroZ_P1_rev | TTGGTCTCAATCTATTAAAGTGCGGCAGATAGAG |
| OMT_P2_fwd | TTGGTCTCAAACCAATGGGCGATACCAAAGAAC |
| OMT_P2_rev | TTGGTCTCATCTTACGGACCTGCTTCGCTAC |
| CAR_P1_fwd | TTGGTCTCATGAAAACAATGACCCAGTCTCACACCC |
| CAR_P1_rev | TTGGTCTCAATCTACAGCAGACCCAGCTGTTTG |
| PPTase_P2_fwd | TTGGTCTCAAACCAATGATCGAAACCATCCTGC |
| PPTase_P2_rev | TTGGTCTCATCTTAAGCGTAAGCGATCGCGG |
| Sfp_P1_fwd | TTGGTCTCATGAAACAATGAAGATTTACGGAATTTATATG |
| Sfp_P1_rev | TTGGTCTCAATCTATAAAAGCTCTTCGTACGAAACCATTG |
| EntD_P2_fwd | TTGGTCTCAAACCAATGGTCGATATGAAAACTAC |
| EntD_P2_rev | TTGGTCTCATCTTAATCGTGTTGGCACAGCG |
| GapC_P1_fwd | TTGGTCTCATGAAAACAATGGCAAAGATAGCTATTAATG |
| GapC_P1_rev | TTGGTCTCAATCTATTTTGCTATTTTTGCAAAGTAAG |
| Pos5c_P2_fwd | TTGGTCTCAAACCAATGAGTACGTTGGATTCACATTCC |
| Pos5c_P2_rev | TTGGTCTCATCTTAATCATTATCAGTCTGTCTC |
| MetK_P1_fwd | TTGGTCTCATGAAAACAATGGCAAAACACCTTTTTACG |
| MetK_OE_rev | CAGCCACGCCTACTGCGTAGGAAACC |
| MetK_OE_fwd | CCTACGCAGTAGGCGTGGCTGAACCG |
| MetK_P1_rev | TTGGTCTCAATCTACTTCAGACCGGCAGCATC |
| HmaS_P1_fwd | TTGGTCTCATGAAAACAATGCAGAATTTCGAAATCG |
| HmaS_P1_rev | TTGGTCTCAATCTAACGCCTCGCGGCTCCAAAC |
| BFD_P1_fwd | TTGGTCTCATGAAAACAATGGCTTCGGTACACGGC |
| BFD_OE_rev | GTCGAACACAGTCTCCGGGTG |
| BFD_OE_fwd | CACCCGGAGACTGTGTTCGAC |
| BFD_P1_rev | TTGGTCTCAATCTACTTCACCGGGCTTACGG |
| HMO_P2_fwd | TTGGTCTCAAACCAATGCGTGAACCGCTGACGC |
| HMO_P2_rev | TTGGTCTCATCTTAGCCGTGAGAACGATCGC |
| HpaB_P1_fwd | TTGGTCTCATGAAAACAATGAAACCCGAAGATTTCCG |
| HpaB_P1_rev | TTGGTCTCAATCTATTGGCGGATGCGATCGAGCAC |
| HpaC_P2_fwd | TTGGTCTCAAACCAATGCAAGTAGATGAACAACG |
| HpaC_P2_rev | TTGGTCTCATCTTAAACAGGCGCTTCCATCTC |
| UbiC_P1_fwd | TTGGTCTCATGAAAACAATGGTCACACCCCGCGTTAAC |
| UbiC_P1_rev | TTGGTCTCAATCTAGGTACAACGGTGACGCCGG |
| PobA_P2_fwd | TTGGTCTCAAACCAATGAAAACTCAGGTTGCAAT |
| PobA_P2_rev | TTGGTCTCATCTTAGGCAACTTCCTCGAACGGC |
| Xfpk_P1_fwd | TTGGTCTCATGAAAACAATGACTTCTCCAGTTATC |
| Xfpk_P1_rev | TTGGTCTCAATCTATTCGTTATCACCAGCAGTTG |
| Pta_P2_fwd | TTGGTCTCAAACCAATGAAGTTGATGGAAAACATC |
| Pta_P2_rev | TTGGTCTCATCTTAACCTTGAGCTTGGGCTTG |

**Table S2** Synthesized genes used in this study.

| **Name** | **Description** |
| --- | --- |
| *OMT* from *H. sapiens* | ATGGGCGATACCAAAGAACAGCGTATTCTGAATCATGTTCTGCAGCATGCCGAACCGGGTAATGCACAGAGCGTTCTGGAAGCAATTGATACCTATTGTGAACAGAAAGAATGGGCCATGAATGTGGGTGATAAAAAAGGCAAAATTGTGGATGCCGTGATCCAAGAACATCAGCCGAGCGTGCTGCTGGAACTGGGTGCATATTGTGGTTATAGCGCAGTTCGTATGGCACGTCTGCTGAGTCCGGGTGCACGTCTGATTACCATTGAAATTAACCCGGATTGTGCAGCAATTACCCAGCGTATGGTTGATTTTGCCGGTGTTAAAGATAAAGTTACCCTGGTTGTTGGTGCAAGCCAGGATATTATTCCGCAGCTGAAAAAAAAATATGACGTGGATACCCTGGATATGGTGTTTCTGGATCATTGGAAAGATCGTTATCTGCCGGATACCCTGCTGCTGGAAGAATGTGGTCTGCTGCGTAAAGGCACCGTTCTGCTGGCAGATAATGTTATTTGTCCTGGTGCACCGGATTTTCTGGCACATGTTCGTGGTAGCAGCTGTTTTGAATGTACCCATTATCAGTCCTTTCTGGAATATCGTGAAGTTGTTGATGGTCTGGAAAAAGCCATCTATAAAGGTCCGGGTAGCGAAGCAGGTCCGTAA |
| *HmaS* from *A. orientalis* | ATGCAGAATTTCGAAATCGACTATGTTGAGATGTATGTGGAGAATCTGGAAGTGGCGGCTTTCTCATGGGTCGACAAGTACGCATTCGCCGTGGCTGGTACGAGCCGTTCAGCGGACCACAGGAGCATCGCGTTGAGACAAGGGCAGGTGACACTTGTTCTTACTGAACCGACGTCAGATAGACATCCGGCCGCCGCGTATTTGCAGACTCATGGCGATGGTGTAGCGGATATTGCTATGGCCACCTCCGACGTTGCTGCGGCCTACGAGGCGGCAGTACGTGCCGGTGCTGAAGCTGTTAGAGCACCAGGGCAACACAGTGAAGCTGCTGTGACGACCGCGACCATAGGTGGGTTTGGAGATGTCGTCCATACTCTGATCCAAAGGGACGGCACTAGCGCTGAATTACCCCCCGGATTTACCGGCTCCATGGACGTTACGAACCATGGTAAAGGAGACGTAGATCTTCTGGGGATAGATCACTTTGCGATATGTCTTAACGCCGGAGATCTGGGACCTACTGTGGAATACTACGAGAGAGCTTTAGGTTTTAGGCAAATATTTGACGAGCATATAGTTGTTGGCGCTCAGGCGATGAACTCAACTGTCGTGCAAAGTGCGAGTGGGGCGGTAACACTAACCTTGATAGAACCCGATCGTAATGCCGACCCCGGGCAAATTGATGAGTTCCTAAAAGATCACCAGGGTGCGGGTGTGCAGCACATCGCCTTTAATTCTAATGACGCTGTGCGTGCAGTGAAAGCACTGTCAGAGAGAGGGGTCGAGTTTTTGAAAACCCCGGGGGCGTATTACGATCTTCTAGGAGAGAGAATAACCCTGCAAACGCATAGTCTGGATGATTTGAGAGCGACCAATGTTTTGGCAGATGAGGATCACGGAGGTCAACTTTTTCAAATCTTCACAGCGAGTACGCACCCAAGACACACCATTTTTTTTGAAGTCATCGAAAGACAAGGAGCGGGCACTTTTGGTTCCAGTAATATCAAGGCTTTATATGAGGCAGTGGAACTAGAACGTACAGGTCAAAGTGAGTTTGGAGCCGCGAGGCGTTAG |
| *HpaB* from *P. aeruginosa* | ATGAAACCCGAAGATTTCCGAGCTTCTGCTACCAGACCATTCACCGGTGAAGAATACTTAGCCAGCCTTCGTGATGATCGTGAAATCTACATCTATGGTGACCGTGTCAAGGACGTTACTTCTCATCCAGCTTTCAGAAACGCTGCTGCTTCTATGGCTAGATTGTACGATGCCTTGCACGACCCACAATCGAAGGAAAAGTTGTGTTGGGAAACTGACACTGGTAACGGTGGTTACACTCACAAGTTCTTCAGATACGCCAGATCCGCTGACGAATTGAGACAACAAAGAGATGCCATCGCTGAATGGTCCAGATTGACCTACGGTTGGATGGGTAGAACCCCAGACTACAAGGCTGCTTTTGGTTCTGCATTAGGTGCTAACCCAGGTTTCTACGGTCGTTTTGAAGACAATGCCAAGACCTGGTATAAGAGAATCCAAGAAGCCTGTTTGTACTTGAACCACGCTATCGTCAACCCACCAATTGACCGTGACAAGCCAGTTGACCAAGTTAAGGATGTTTTCATCTCTGTCGATGAAGAAGTCGATGGTGGTATTGTCGTCAGTGGTGCTAAGGTTGTTGCTACCAACTCCGCTCTAACCCACTACAACTTCGTTGGTCAAGGTTCTGCTCAATTGTTGGGTGACAACACCGATTTCGCTTTGATGTTCATTGCTCCAATGAACACTCCAGGTATGAAGTTGATCTGTCGGCCTTCTTACGAATTGGTCGCTGGTATCGCCGGTTCTCCATTTGACTACCCTTTGTCTTCTAGATTCGATGAAAACGACGCTATTTTGGTTATGGACAAGGTGTTCATTCCTTGGGAAAACGTTTTGATTTACAGAGACTTCGAAAGATGTAAGCAATGGTTTCCACAAGGTGGTTTCGGCAGATTATTCCCAATGCAAGGTTGTACTAGATTAGCTGTCAAATTGGACTTCATCACTGGTGCCCTTTACAAGGCTTTGCAATGCACCGGTTCCTTGGAATTTCGTGGTGTCCAAGCTCAAGTCGGTGAAGTTGTTGCTTGGAGAAACTTGTTCTGGTCTTTGACAGATGCTATGTACGGTAACGCGTCTGAATGGCACGGTGGTGCTTTCTTGCCATCTGCCGAGGCTTTGCAAGCCTACAGAGTTTTGGCTCCACAAGCCTACCCAGAAATCAAGAAGACTATTGAACAAGTTGTCGCTTCTGGTTTGATTTACTTGCCATCCGGTGTTAGAGACTTACACAACCCTCAACTGGACAAGTACCTATCCACTTACTGTAGAGGTTCTGGAGGTATGGGTCACAGAGAAAGAATCAAGATTTTGAAGCTATTGTGGGATGCTATCGGGTCAGAATTTGGTGGTAGACACGAATTATACGAAATCAACTACGCCGGTTCCCAGGATGAAATAAGAATGCAAGCCTTGAGACAAGCCATTGGTTCCGGTGCCATGAAGGGTATGTTGGGTATGGTTGAACAATGTATGGGTGATTATGACGAAAATGGTTGGACTGTTCCACATTTGCATAACCCAGATGACATTAACGTGCTCGATCGCATCCGCCAATGA |
| *Xfpk* from *B. breve* | ATGACTTCTCCAGTTATCGGTACTCCTTGGAAGAAGTTGAACGCTCCTGTTTCAGAAGAATCTTTGGAAGGTGTTGACAAGTACTGGAGAGTTGCTAACTACTTGTCCATCGGTCAAATCTACTTAAGATCCAACCCATTGATGAAGGCCCCATTCACTAGAGAAGATGTTAAGCACCGTTTGGTTGGTCACTGGGGTACTACCCCAGGTTTGAACTTCTTGATTGGTCACATCAACAGATTCATCGCTGACCACGGTCAAAACACCGTCATCATTATGGGTCCAGGTCACGGTGGACCGGCTGGTACTTCTCAGTCCTATTTGGATGGTACTTACACCGAAACTTTCCCAAAAATCACCAAGGACGAAGCTGGCTTGCAAAAGTTTTTCAGACAATTCTCCTACCCAGGTGGTATTCCATCTCATTTCGCTCCAGAAACTCCTGGTTCTATCCACGAAGGTGGTGAATTGGGTTACGCCTTGTCCCACGCGTACGGTGCTATCATGGACAACCCCTCTTTATTCGTTCCAGCTATTGTCGGTGATGGCGAAGCCGAAACTGGTCCATTGGCTACCGGTTGGCAATCTAACAAATTGGTGAACCCACGTACCGATGGTATTGTTTTGCCAATTTTGCATTTGAACGGTTATAAGATAGCTAACCCAACCATTCTGTCTAGAATTTCCGATGAAGAGTTGCACGAATTTTTCCACGGTATGGGTTACGAACCATACGAATTTGTTGCTGGTTTCGACGATGAAGATCACATGTCTATTCACAGAAGATTTGCTGAATTGTGGGAAACCATCTGGGACGAAATTTGCGACATCAAGGCGGCTGCTCAAACTGACAACGTCCACAGACCATTCTACCCAATGTTGATCTTCAGAACCCCAAAGGGTTGGACTTGTCCAAAGTACATTGACGGCAAGAAGACTGAAGGTTCTTGGAGAGCCCACCAAGTTCCACTCGCTTCGGCTAGAGACACTGAAGCCCACTTCGAAGTTCTGAAGAACTGGTTAGAATCCTACAAGCCAGAAGAATTATTTGATGCCAACGGGGCAGTCAAGGATGACGTCTTGGCCTTCATGCCAAAGGGTGAATTGAGAATTGGTGCTAACCCAAATGCTAATGGTGGTGTTATTAGGGATGATTTGAAGCTGCCAAACTTGGAAGACTACGAAGTCAAGGAAGTTGCTGAATACGGTCACGGCTGGGGTCAATTGGAAGCCACCCGTACGCTAGGTGCTTATACCCGGGATATAATCAGAAACAACCCTAGAGATTTCAGAATTTTCGGTCCAGACGAAACTGCCTCCAACAGATTGCAAGCCTCTTACGAAGTAACAAACAAGCAATGGGATGCTGGTTACATCTCTGACGAAGTCGATGAACACATGCATGTTTCCGGTCAAGTTGTTGAACAATTGTCTGAACATCAAATGGAAGGTTTTTTGGAAGCCTACTTGCTAACTGGTAGACACGGTATCTGGTCTTCTTACGAATCTTTCGTTCATGTCATCGACTCCATGTTGAACCAACACGCTAAGTGGTTGGAAGCTACTGTCAGAGAAATCCCTTGGAGAAAGCCAATCGCTTCGATGAACTTGTTGGTTTCTTCTCACGTTTGGCGTCAAGATCATAACGGTTTCTCTCACCAAGACCCAGGTGTTACTAGCGTTCTATTGAATAAGTGTTTCCACAACGATCACGTTATCGGTATTTACTTCGCCACCGACGCTAACATGTTGTTGGCTATCGCTGAAAAGTGTTACAAGTCTACCAACAAGATCAACGCCATTATTGCCGGTAAGCAACCAGCCGCTACCTGGTTAACTTTGGACGAAGCCAGAGCTGAACTGGCCAAGGGTGCTGCAGCTTGGGACTGGGCTTCCACTGCCAAGAACAATGATGAAGCTGAGGTCGTCTTGGCTGCCGCTGGTGACGTCCCAACTCAAGAAATTATGGCTGCGTCTGACAAGTTGAAGGAATTAGGTGTCAAATTCAAGGTTGTTAACGTTGCTGACTTGTTATCTTTGCAAAGTGCTAAGGAAAACGATGAAGCCTTGTCTGACGAAGAATTTGCTGATATTTTCACTGCTGACAAACCAGTCTTGTTCGCTTACCACTCTTACGCTCACGACGTTAGAGGTTTGATTTACGACAGACCAAACCATGATAACTTCAACGTTCACGGTTACGAAGAAGAAGGTTCCACCACCACCCCATACGACATGGTTAGAGTCAATAGAATCGATAGATACGAATTAACCGCTGAGGCTTTGAGAATGATCGATGCTGATAAGTACGCTGATAAGATAGACGAATTAGAAAAGTTCCGTGACGAAGCCTTCCAATTTGCTGTTGACAAAGGTTACGACCACCCAGATTACACAGACTGGGTTTACTCTGGTGTTAACACTGACAAGAAAGGTGCTGTCACTGCCACTGCCGCAACTGCTGGTGATAACGAATAG |
| *Pta* from *C. kluyveri* | ATGAAGTTGATGGAAAACATCTTCGGACTGGCTAAGGCTGACAAGAAGAAGATTGTCTTAGCTGAAGGTGAAGAAGAGCGTAACATTCGTGCTTCCGAAGAAATCATCAGAGATGGTATTGCTGACATTATCTTGGTCGGTTCTGAATCCGTTATCAAGGAAAACGCCGCTAAGTTCGGTGTTAACTTGGCTGGTGTCGAAATTGTTGACCCAGAAACTTCTTCTAAGACCGCCGGTTATGCTAACGCTTTCTACGAAATTAGAAAGAACAAGGGTGTTACTTTGGAAAAGGCTGATAAGATCGTTAGAGATCCTATCTACTTCGCTACCATGATGGTTAAGTTGGGTGACGCTGATGGTTTGGTTTCCGGTGCCATTCACACTACTGGTGATTTGTTAAGACCAGGCTTGCAAATCGTCAAGACTGTCCCAGGTGCTAGTGTTGTCTCTTCCGTTTTCTTAATGTCTGTTCCAGATTGTGAATACGGTGAAGATGGTTTCTTGTTATTTGCCGACTGTGCTGTCAACGTCTGTCCAACTGCTGAAGAATTATCTTCAATTGCCATCACCACTGCCGAAACCGCCAAGAACTTGTGTAAGATTGAACCAAGAGTTGCTATGTTGTCTTTCTCTACTATGGGTTCTGCTTCCCACGAATTGGTTGACAAAGTTACCAAGGCCACCAAGTTGGCTAAAGAAGCTAGACCAGACTTGGACATTGACGGTGAACTACAATTGGATGCTTCTTTGGTCAAAAAAGTGGCAGACTTGAAGGCTCCAGGTTCCAAGGTTGCTGGTAAGGCTAATGTTTTGATTTTCCCAGACATCCAAGCTGGTAACATTGGTTACAAGCTAGTTCAAAGATTCGCCAAAGCTGAAGCTATCGGTCCAATCTGCCAAGGTTTTGCTAAGCCAATCAACGATTTGAGCAGAGGTTGTTCTGTCGATGACATCGTCAAGGTTGTTGCTGTCACCGCCGTCCAAGCCCAAGCTCAAGGTTAA |

**Table S3** Plasmids used in this study.

| **Name** | **Description** |
| --- | --- |
| p414-TEF1-Cas9 ^1^ | Plasmid harboring *Cas9* gene under the control of TEF1 promoter with TRP selection marker |
| pRS426SNR52 | Plasmid harboring P_SNR52_-T_SUP4_ cassette, lab stock |
| pRS426-gRNA(adh6) ^2^ | pRS426SNR52 derivative with P_SNR52_-gRNA_adh6-T_SUP4_ |
| pRS426-gRNA(adh7) ^2^ | pRS426SNR52 derivative with P_SNR52_-gRNA_adh7-T_SUP4_ |
| pRS426-gRNA(sfa1) ^2^ | pRS426SNR52 derivative with P_SNR52_-gRNA_sfa1-T_SUP4_ |
| pRS426-gRNA(gre2) ^2^ | pRS426SNR52 derivative with P_SNR52_-gRNA_gre2-T_SUP4_ |
| pRS426-gRNA(hfd1) ^2^ | pRS426SNR52 derivative with P_SNR52_-gRNA_hfd1-T_SUP4_ |
| pRS426-gRNA(gre3) | pRS426SNR52 derivative with P_SNR52_-gRNA_gre3-T_SUP4_ |
| pRS426-gRNA(ydl124w) | pRS426SNR52 derivative with P_SNR52_-gRNA_ydl124w-T_SUP4_ |
| pRS426-gRNA(gcy1) | pRS426SNR52 derivative with P_SNR52_-gRNA_gcy1-T_SUP4_ |
| pRS426-gRNA(ypr1) | pRS426SNR52 derivative with P_SNR52_-gRNA_ypr1-T_SUP4_ |
| pRS426-gRNA(ari1) | pRS426SNR52 derivative with P_SNR52_-gRNA_ari1-T_SUP4_ |
| pRS426-gRNA(ydr541c) | pRS426SNR52 derivative with P_SNR52_-gRNA_ydr541c-T_SUP4_ |
| pRS426-gRNA(aad3) | pRS426SNR52 derivative with P_SNR52_-gRNA_aad3-T_SUP4_ |
| pRS426-gRNA(bts1) ^2^ | pRS426SNR52 derivative with P_SNR52_-gRNA_bts1-T_SUP4_ |
| pRS426-gRNA(ypl062w) ^2^ | pRS426SNR52 derivative with P_SNR52_-gRNA_ypl062w-T_SUP4_ |
| pRS426-gRNA(rox1) ^2^ | pRS426SNR52 derivative with P_SNR52_-gRNA_rox1-T_SUP4_ |
| pRS426-gRNA(yjl064w) ^2^ | pRS426SNR52 derivative with P_SNR52_-gRNA_yjl064w-T_SUP4_ |
| pRS426-gRNA(pdc6) | pRS426SNR52 derivative with P_SNR52_-gRNA_pdc6-T_SUP4_ |
| pRS426-gRNA(pha2) | pRS426SNR52 derivative with P_SNR52_-gRNA_pha2-T_SUP4_ |
| pRS426-gRNA(are1) | pRS426SNR52 derivative with P_SNR52_-gRNA_are1-T_SUP4_ |
| pRS426-gRNA(gdh1) | pRS426SNR52 derivative with P_SNR52_-gRNA_gdh1-T_SUP4_ |
| pRS426-gRNA(gpp1) | pRS426SNR52 derivative with P_SNR52_-gRNA_gpp1-T_SUP4_ |
| pRS426-gRNA(gal80N) | pRS426SNR52 derivative with P_SNR52_-gRNA_gal80N-T_SUP4_ |
| pRS423-AsbF/OMT | pRS423A-GGA derivative with P_GAL10_-*AsbF*-T_ADH1_; P_GAL1_-*OMT*-T_CYC1_ |
| pRS423-AroZ/OMT | pRS423A-GGA derivative with P_GAL10_-*AroZ*-T_ADH1_; P_GAL1_-*OMT*-T_CYC1_ |
| pRS425-Sfp/EntD | pRS425A-GGA derivative with P_GAL10_-*Sfp*-T_ADH1_; P_GAL1_-*EntD*-T_CYC1_ |
| pRS423-MetK/OMT | pRS423A-GGA derivative with P_GAL10_-*MetK*-T_ADH1_; P_GAL1_-*OMT*-T_CYC1_ |
| pRS423-MetK^I303V^/OMT | pRS423A-GGA derivative with P_GAL10_-*MetK^I303V^*-T_ADH1_; P_GAL1_-*OMT*-T_CYC1_ |
| pRS425-GapC/Pos5c | pRS425A-GGA derivative with P_GAL10_-*GapC*-T_ADH1_; P_GAL1_-*Pos5c*-T_CYC1_ |
| pRS425-CAR/PPTase | pRS425A-GGA derivative with P_GAL10_-*CAR*-T_ADH1_; P_GAL1_-*PPTase*-T_CYC1_ |
| pRS423-HpaB/HpaC | pRS423A-GGA derivative with P_GAL10_-*HpaB*-T_ADH1_; P_GAL1_-*HpaC*-T_CYC1_ |
| pRS425-BFD/HMO | pRS425A-GGA derivative with P_GAL10_-*BFD*-T_ADH1_; P_GAL1_-*HMO*-T_CYC1_ |
| pRS426-HmaS/OMT | pRS426A-GGA derivative with P_GAL10_-*HmaS*-T_ADH1_; P_GAL1_-*OMT*-T_CYC1_ |
| pRS426-UbiC/PobA | pRS426A-GGA derivative with P_GAL10_-*UbiC*-T_ADH1_; P_GAL1_-*PobA*-T_CYC1_ |
| pRS423-Xfpk/Pta | pRS423A-GGA derivative with P_GAL10_-*Xfpk*-T_ADH1_; P_GAL1_-*Pta*-T_CYC1_ |

**Table S4** Strains used in this study.

| **Name** | **Description** |
| --- | --- |
| JS-CR(2M) ^3^ | Strain BY4741 derivative with Δ*aro10*::P_GAL10_-*Aro4^fbr^*-T_CYC1_ and P_GAL1_-*Aro8*-T_CYC1_; Δ*trp1*:: P_GAL10_-*Aro7^fbr^*-T_CYC1_ and P_GAL1_-*Aro8*-T_CYC1_; P_CUP1_-*Gal4*; P_CTR1_-*Gal80* |
| JS-RARE1 | Strain JS-CR(2M) derivative with ∆*adh6*, ∆*adh7*, ∆*sfa1*, ∆*gre2* and ∆*hfd1* |
| JS-RARE2 | Strain JS-RARE1 derivative with ∆*gre3*, ∆*ydl124w*, ∆*gcy1*, and ∆*ypr1* |
| JS-RARE3 | Strain JS-RARE2 derivative with ∆*ari1*, ∆*ydr541c*, and ∆*aad3* |
| JS-CR(2M)-P1 | Strain JS-CR(2M) transformed with pRS423-AroZ/OMT and pRS425-CAR/PPTase |
| JS-RARE3-P2 | Strain JS-RARE3 transformed with pRS423-AroZ/OMT and pRS425-CAR/PPTase |
| JS-RARE3-P3 | Strain JS-RARE3 transformed with pRS423-HpaB/HpaC, pRS425-BFD/HMO, and pRS426-HmaS/OMT |
| JS-RARE3-P4 | Strain JS-RARE3 transformed with pRS425-CAR/PPTase and pRS426-UbiC/PobA |
| JS-VA1a | Strain JS-RARE3 derivative with Δ*rox1*::P_GAL10_-*AroZ*-T_ADH1_; P_GAL1_-*OMT*-T_CYC1_, Δ*bts1*::P_GAL10_-*CAR*-T_ADH1_; P_GAL1_-*PPTase*-T_CYC1_ |
| JS-VA1b | Strain JS-RARE3 derivative with Δ*rox1*::P_GAL10_-*AsbF*-T_ADH1_; P_GAL1_-*OMT*-T_CYC1_, Δ*bts1*::P_GAL10_-*CAR*-T_ADH1_; P_GAL1_-*PPTase*-T_CYC1_ |
| JS-VA2 | Strain JS-RARE3 derivative with Δ*rox1*::P_GAL10_-*AroZ*-T_ADH1_; P_GAL1_-*OMT*-T_CYC1_, Δ*bts1*::P_GAL10_-*CAR*-T_ADH1_; P_GAL1_-*PPTase*-T_CYC1_, Δ*ypl062w*::P_GAL10_-*Sfp*-T_ADH1_; P_GAL1_-*EntD*-T_CYC1_ |
| JS-VA3 | Strain JS-RARE3 derivative with Δ*rox1*::P_GAL10_-*AroZ*-T_ADH1_; P_GAL1_-*OMT*-T_CYC1_, Δ*bts1*::P_GAL10_-*CAR*-T_ADH1_; P_GAL1_-*PPTase*-T_CYC1_, Δ*ypl062w*::P_GAL10_-*Sfp*-T_ADH1_; P_GAL1_-*EntD*-T_CYC1_, Δ*yjl064w*::P_GAL10_-*GapC*-T_ADH1_; P_GAL1_-*Pos5c*-T_CYC1_ |
| JS-VA4 | Strain JS-RARE3 derivative with Δ*rox1*::P_GAL10_-*AroZ*-T_ADH1_; P_GAL1_-*OMT*-T_CYC1_, Δ*bts1*::P_GAL10_-*CAR*-T_ADH1_; P_GAL1_-*PPTase*-T_CYC1_, Δ*ypl062w*::P_GAL10_-*Sfp*-T_ADH1_; P_GAL1_-*EntD*-T_CYC1_, Δ*yjl064w*::P_GAL10_-*MetK*-T_ADH1_; P_GAL1_-*OMT*-T_CYC1_ |
| JS-VA5 | Strain JS-RARE3 derivative with Δ*rox1*::P_GAL10_-*AroZ*-T_ADH1_; P_GAL1_-*OMT*-T_CYC1_, Δ*bts1*::P_GAL10_-*CAR*-T_ADH1_; P_GAL1_-*PPTase*-T_CYC1_, Δ*ypl062w*::P_GAL10_-*Sfp*-T_ADH1_; P_GAL1_-*EntD*-T_CYC1_, Δ*yjl064w*::P_GAL10_-*MetK^I303V^*-T_ADH1_; P_GAL1_-*OMT*-T_CYC1_ |
| JS-VA6 | Strain JS-RARE3 derivative with Δ*rox1*::P_GAL10_-*AroZ*-T_ADH1_; P_GAL1_-*OMT*-T_CYC1_, Δ*bts1*::P_GAL10_-*CAR*-T_ADH1_; P_GAL1_-*PPTase*-T_CYC1_, Δ*ypl062w*::P_GAL10_-*Sfp*-T_ADH1_; P_GAL1_-*EntD*-T_CYC1_, Δ*yjl064w*::P_GAL10_-*MetK^I303V^*-T_ADH1_; P_GAL1_-*OMT*-T_CYC1_, Δ*pdc6*::P_GAL10_-*GapC* -T_ADH1_; P_GAL1_-*Pos5c*-T_CYC1_ |
| JS-VA7 | Strain JS-RARE3 derivative with Δ*rox1*::P_GAL10_-*AroZ*-T_ADH1_; P_GAL1_-*OMT*-T_CYC1_, Δ*bts1*::P_GAL10_-*CAR*-T_ADH1_; P_GAL1_-*PPTase*-T_CYC1_, Δ*ypl062w*::P_GAL10_-*Sfp*-T_ADH1_; P_GAL1_-*EntD*-T_CYC1_, Δ*yjl064w*::P_GAL10_-*MetK^I303V^*-T_ADH1_; P_GAL1_-*OMT*-T_CYC1_, Δ*pdc6*::P_GAL10_-*GapC* -T_ADH1_; P_GAL1_-*Pos5c*-T_CYC1_, Δ*pha2*::P_GAL10_-*HmaS*-T_ADH1_; P_GAL1_-*OMT*-T_CYC1_, Δ*are1*::P_GAL10_-*BFD*-T_ADH1_; P_GAL1_-*HMO*-T_CYC1_, Δ*gdh1*::P_GAL10_-*HpaB*-T_ADH1_; P_GAL1_-*HpaC*-T_CYC1_ |
| JS-VA8 | Strain JS-RARE3 derivative with Δ*rox1*::P_GAL10_-*AroZ*-T_ADH1_; P_GAL1_-*OMT*-T_CYC1_, Δ*bts1*::P_GAL10_-*CAR*-T_ADH1_; P_GAL1_-*PPTase*-T_CYC1_, Δ*ypl062w*::P_GAL10_-*Sfp*-T_ADH1_; P_GAL1_-*EntD*-T_CYC1_, Δ*yjl064w*::P_GAL10_-*MetK^I303V^*-T_ADH1_; P_GAL1_-*OMT*-T_CYC1_, Δ*pdc6*::P_GAL10_-*GapC* -T_ADH1_; P_GAL1_-*Pos5c*-T_CYC1_, Δ*gdh1*::P_GAL10_-*UbiC* -T_ADH1_; P_GAL1_-*PobA*-T_CYC1_ |
| JS-VA9 | Strain JS-RARE3 derivative with Δ*rox1*::P_GAL10_-*AroZ*-T_ADH1_; P_GAL1_-*OMT*-T_CYC1_, Δ*bts1*::P_GAL10_-*CAR*-T_ADH1_; P_GAL1_-*PPTase*-T_CYC1_, Δ*ypl062w*::P_GAL10_-*Sfp*-T_ADH1_; P_GAL1_-*EntD*-T_CYC1_, Δ*yjl064w*::P_GAL10_-*MetK^I303V^*-T_ADH1_; P_GAL1_-*OMT*-T_CYC1_, Δ*pdc6*::P_GAL10_-*GapC* -T_ADH1_; P_GAL1_-*Pos5c*-T_CYC1_, Δ*gdh1*::P_GAL10_-*UbiC* -T_ADH1_; P_GAL1_-*PobA*-T_CYC1_, Δ*gpp1*::P_GAL10_-*Xfpk*-T_ADH1_; P_GAL1_-*Pta*-T_CYC1_ |
| JS-VA10 | Strain JS-RARE3 derivative with Δ*rox1*::P_GAL10_-*AroZ*-T_ADH1_; P_GAL1_-*OMT*-T_CYC1_, Δ*bts1*::P_GAL10_-*CAR*-T_ADH1_; P_GAL1_-*PPTase*-T_CYC1_, Δ*ypl062w*::P_GAL10_-*Sfp*-T_ADH1_; P_GAL1_-*EntD*-T_CYC1_, Δ*yjl064w*::P_GAL10_-*MetK^I303V^*-T_ADH1_; P_GAL1_-*OMT*-T_CYC1_, Δ*pdc6*::P_GAL10_-*GapC* -T_ADH1_; P_GAL1_-*Pos5c*-T_CYC1_, Δ*gdh1*::P_GAL10_-*UbiC*-T_ADH1_; P_GAL1_-*PobA*-T_CYC1_, Δ*gpp1*::P_GAL10_-*Xfpk*-T_ADH1_; P_GAL1_-*Pta*-T_CYC1_, ΔP*_GAL80_ -GAL80*::P*_CTR1_*-*K15*∼*GAL80* |
